# α-Copaene is a potent repellent against the Asian Citrus Psyllid *Diaphorina citri*

**DOI:** 10.1101/2024.10.04.616677

**Authors:** Rodrigo Facchini Magnani, Haroldo Xavier Linhares Volpe, Rejane Angélica Grigio Luvizotto, Tatiana Aparecida Mulinari, Thiago Trevisoli Agostini, Jairo Kenupp Bastos, Victor Pena Ribeiro, Michele Carmo-Sousa, Nelson Arno Wulff, Leandro Peña, Walter S. Leal

## Abstract

The Asian Citrus Psyllid (ACP), *Diaphorina citri*, severely threatens citrus production worldwide by transmitting the greening(= Huanglongbing)-causing bacterium *Candidatus* Liberibacter asiaticus. There is growing evidence that the push-pull strategy is suitable to partially mitigate HLB by repelling ACP with transgenic plants engineered to produce repellents and attracting the vector to plants with a minimal disease transmission rate. Species that pull ACP away from commercial citrus plants have been identified, and transgenic plants that repel ACP have been developed. The concept of a repellent-producing plant was first demonstrated with an *Arabidopsis* line engineered to overexpress a gene controlling the synthesis of β-caryophyllene and other sesquiterpenes. We have analyzed the volatile organic compounds released by this *Arabidopsis* line and identified α-humulene, α-copaene, and trace amounts of β-elemene, in addition to β-caryophyllene. Behavioral measurements demonstrated that α-copaene repels ACP at doses ca. 100x lower than those needed for β-caryophyllene repellence. In contrast, α-humulene is innocuous at the level emitted by the transgenic plant. We confirmed that a mixture of the three sesquiterpenes in the ratio 100:10:1 repels ACP. Likewise, a commercial sample of refined copaiba oil containing the three sesquiterpenes, in a proportion similar to that in the transgenic plant, repelled ACP.

## Introduction

The Asian Citrus Psyllid (ACP), *Diaphorina citri* (Hemiptera: Psyllidae), is a vector of the bacterium *Candidatus* Liberibacter asiaticus (CLas), which causes the citrus disease known as huanglongbing (HLB) or greening. ^1^ HLB has already decimated the citrus industry in Florida and continues to be a severe threat to citrus production worldwide. Brazil is the largest orange producer in the world and has been sustaining severe losses since 2004 when HLB was detected for the first time ^2^ in the State of São Paulo. The latest assessment indicates a greening incidence rate of 44.35% (90.6 million citrus trees) in the State of São Paulo and the State of Minas Gerais West-Southwest region. ^3^ Therefore, strategies to control ACP populations and/or reduce *C*Las transmission are sorely needed. One promising environmentally sustainable approach to mitigate ACP’s threat is push-pull, ^4^ which integrates lures to attract (pull) and stimuli to repel (push) insect pests and disease vectors. For example, ornamental and other Rutaceae plants that attract ACP to a refuge area may be combined with transgenic plants that produce ACP repellents. ^5^ Orange jasmine, *Murraya paniculata* (Rutaceae), known in Brazil as murta, has been demonstrated to attract ACP, ^6^ but only 1% of psyllids may be infected by feeding on this plant species. ^7^ On the other hand, the concept of plants that repel ACP was first shown with a transgenic *Arabidopsis* line ^8^ and, subsequently, with sweet orange plants. ^9^

The first ACP-repellent transgenic *Arabidopsis* line ^8^ was engineered to overexpress the sesquiterpene synthase gene At5g44630, which was previously demonstrated to be involved in the synthesis of β-caryophyllene, α-humulene, α-copaene, and β-elemene. ^10^ Biosynthetically, these sesquiterpenes are derived from farnesyl pyrophosphate (FPP) by distinct cyclization and derivatization pathways. For example, while β-caryophyllene is derived from the cyclization through FPP’s carbons 1 and 11, α-copaene is produced after the cyclization of FPP’s carbons 1 and 10. Thus, the *Arabidopsis* line engineered to produce β-caryophyllene ^8^ is expected to make other sesquiterpenes, although we cannot predict their ratios. Here, we report that the *Arabidopsis* line yields β-caryophyllene, α-humulene, and α-copaene in the nominal ratio of 100:10:1, and trace amounts of β-elemene. Behavioral measurements demonstrated that α-copaene is a potent repellent effective at 100x lower doses than β-caryophyllene. Although these two sesquiterpene repellents did not synergize, when combined, they seemed to overload the ACP’s olfactory system, thus requiring a lower dose to repel ACP. Behavioral measurements suggest that α-humulene is innocuous at the levels produced by the transgenic plant. Therefore, the tertiary mixture emitted by the transgenic plant effectively repels ACP. Lastly, we demonstrate that a commercially available refined copaiba oil containing β-caryophyllene, α-humulene, and α-copaene in a ratio similar to that yielded by the transgenic plant is also an effective ACP repellent.

## Results and Discussion

### The ratio of caryophyllene isomers produced by a transgenic *Arabidopsis* line

Previously, we have demonstrated that ACP is repelled by authentic *β*-caryophyllene and volatiles emitted by *A. thaliana* line overexpressing synthase At5g23960. ^8^ Because β-caryophyllene was also the main volatile organic compound (VOC) emitted by guava plants (*Psidium guajava*) known to repel ACP, ^11^ it was concluded that this sesquiterpene was the main active ingredient in the transgenic *Arabidopsis* line.

Considering that in *A. thaliana*, At5g23960 converts farnesyl pyrophosphate (FPP) into β-caryophyllene, α-copaene, α-humulene, and β-elemene ^10,12^ by different FPP cyclization pathways, we analyzed VOCs to determine the ratio of these sesquiterpenes in the transgenic *Arabidopsis* line.

Thermal desorber–gas chromatography-mass spectrometry (TD-GC-MS) analyses identified *β*-caryophyllene, α-copaene, and α-humulene in the VOCs emitted by the transgenic *Arabidopsis* line (Fig. 1) based on their retention times, mass spectra, and NIST (National Institute of Standards and Technology) library searches. Analyses of authentic standards of β-caryophyllene, α-copaene, and α-humulene corroborated these initial identifications. Given that the absolute configurations of *A. thaliana-*derived *caryophyllene* and copaene are already known ^10^ as (-)-(*E*)-caryophyllene (=β-caryophyllene, IUPAC name, (1*R*,4*E*,9*S*)-4,11,11-trimethyl-8-methylidenebicyclo[7.2.0]undec-4-ene, CAS#87-44-5) and (-)-α-copaene (=copaene, IUPAC name, (1*R*,2*S*,6*S*,7*S*,8*S*)-1,3-dimethyl-8-propan-2-yltricyclo[4.4.0.0^2,7^]dec-3-ene, CAS#3856-25-5), we did not pursue stereochemistry analysis. Likewise, humulene from *A. thaliana* has been identified ^10^ as α-humulene (=humulene, α-caryophyllene, IUPAC name, (1*E*,4*E*,8*E*)-2,6,6,9-tetramethylcycloundeca-1,4,8-triene, CAS# 6753-98-6). The transgenic *Arabidopsis* line VOC samples also contained trace amounts of β-elemene.

**Figure 1.**
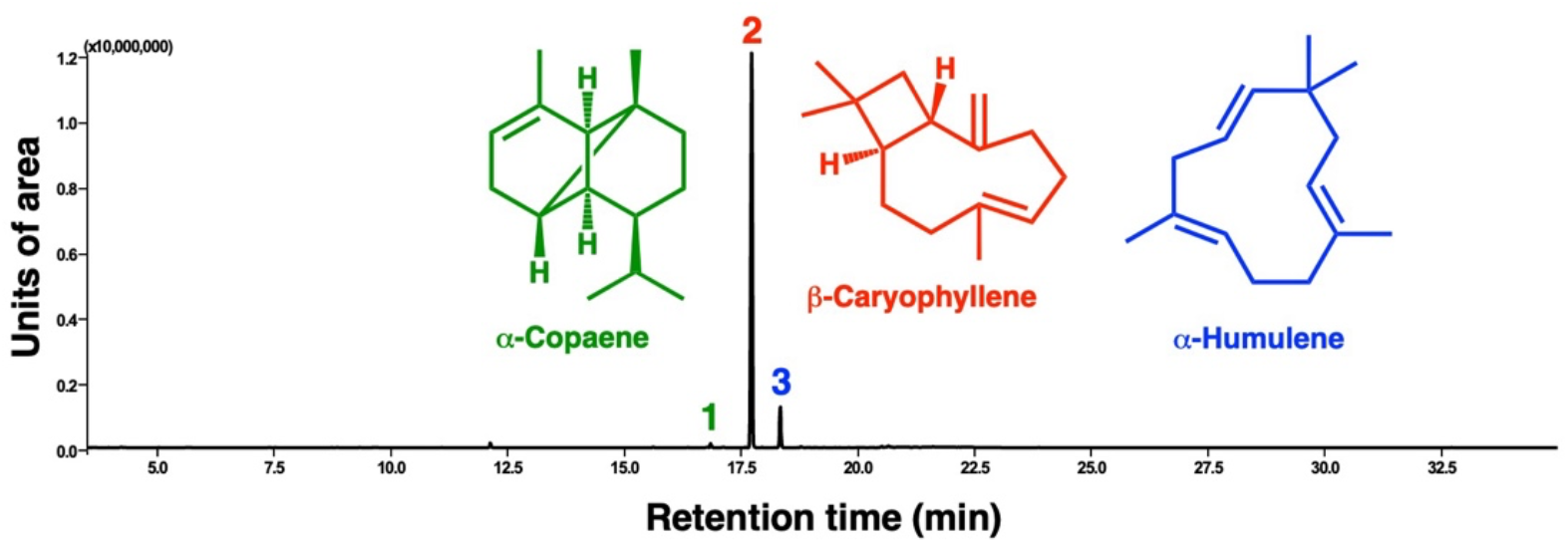
Chromatogram of volatiles emitted by an overexpression line of *Arabidopsis thaliana*. TD-GC-MS analyzed VOCs emitted by the transgenic line, which were captured by Tenax^®^ TA columns, desorbed, cryogenically recaptured, and injected into a gas chromatograph coupled with a mass spectrometer. This representative total ion monitoring trace shows three prominent peaks, which were identified as (1) α-copaene, (2) β-caryophyllene, and (3) α-humulene.

With seven replicates, we determine the relative ratio of the three sesquiterpenes in airborne volatiles emitted by the transgenic *Arabidopsis* line to be 0.95 ± 0.06/100/9.82 ± 0.44 (α-copaene, β-caryophyllene and α-humulene). In sum, β-caryophyllene is released on average 10x more than α-humulene and 100x more than α-copaene.

For behavioral studies involving mixtures of semiochemicals with different vapor pressures (like emanations from the transgenic *Arabidopsis* line), it is crucial to use devices that faithfully release semiochemicals in the same ratios as their sources and in a steady fashion. ^13,14^

### Adjusting β-caryophyllene concentrations released by static and dynamic devices

For subsequent behavioral measurements, we used a dynamic device inspired by the “wick-baits” ^14^ designed to release a steady flow of semiochemical mixtures faithfully representing the proportion of the components in a mixture. ^13^ We have already demonstrated that β-caryophyllene at 1 µg/µl released from a static device (open glass vial) elicited repellence activity in ACP. ^8^ First, we determined the concentration of β-caryophyllene in our dynamic device equivalent to the 1 µg/µl dose in the static device. For that, we captured β-caryophyllene from the static device (n = 17) and generated a calibration curve using 0.05, 0.1, 0.3, 0.5, and 0.8 µg/µl of β-caryophyllene solutions loaded into the dynamic devices (n = 3 for each dose). These analyses showed that the amount of β-caryophyllene in the airborne volatiles emitted from a static device (source dose, 1 µg/µl dose) is equivalent to 0.17 ± 0.01 µg/µl (mean ± sem; throughout the paper means are accompanied by the standard error of the means).

Next, we measured ACP behavioral responses to β-caryophyllene (0.17 µg/µl source dose) released from our dynamic device. ACP females spend significantly more time in the control (hexane) than in the β-caryophyllene field (n = 99, p = 0.0152, Wilcoxon matched-pairs signed-ranked test; hereafter Wilcoxon test; mean residence times in the β-caryophyllene and control fields were 4.06 ± 0.39 min and 5.94 ± 0.39 min, respectively) (Fig. 2A). There were no significant differences between treatment and control when tested at lower and higher doses of β-caryophyllene (0.13 µg/µl, n = 103, p = 0.3792; 0.22 µg/µl, n = 111, p = 0.1569) (Fig. 2B,C).

**Figure 2.**
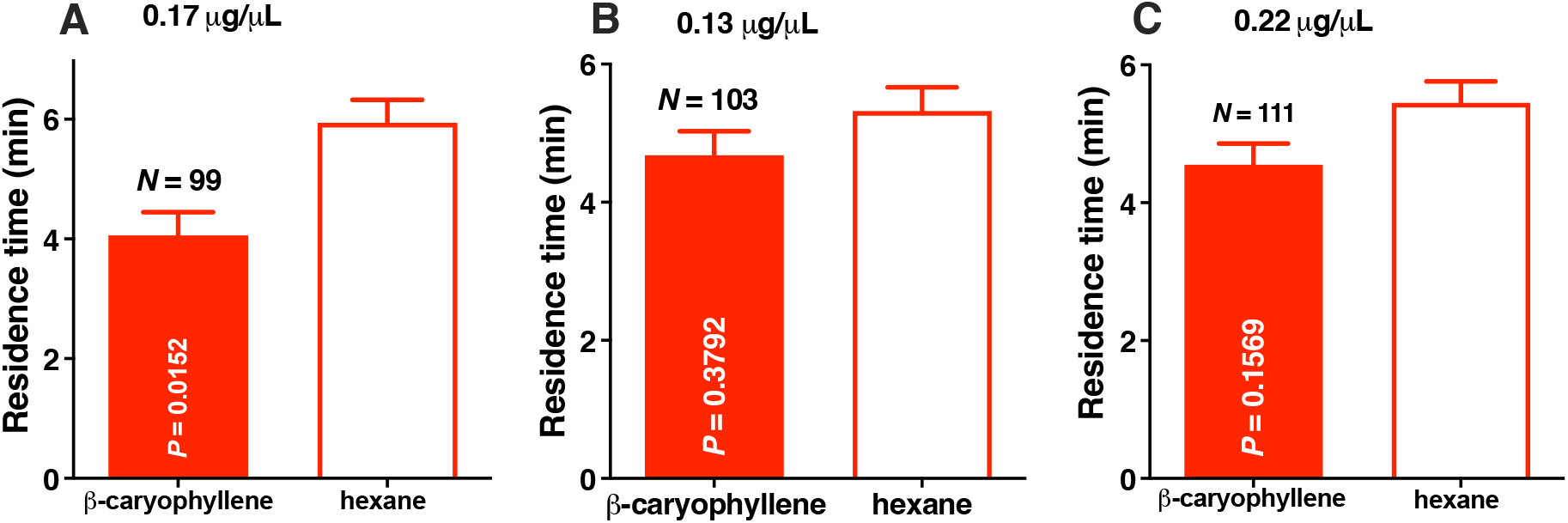
ACP responses to β-caryophyllene released from a dynamic device. (A) ACP females were repelled by β-caryophyllene released from a wick-based dynamic device at 0.17 µg/µl as demonstrated by the significantly longer residence time in the control than in the treatment fields (n = 99, p = 0.0152, Wilcoxon test). By contrast, ACP females were not repelled by β-caryophyllene (B) at lower (0.13 µg/µl, n = 103, p = 0.3792) or (C) higher (0.22 µg/µl, n = 111, p = 0.1569) doses.

Next, we measured the rate of β-caryophyllene emitted from our dynamic device loaded with the active dose of 0.17 µg/µl. VOCs released from the device were captured by Tenax^®^ TA columns and analyzed by TD-GC-MS. The release rate of β-caryophyllene was 0.044 ± 0.013 µg/min (n = 21).

Additionally, we measured the time-release relationships of β-caryophyllene, α-humulene, and α-copaene to determine the stability of our dynamic device throughout repellency assays (15 min; behavioral measurement time, 10 min). Solutions (n = 11) of authentic β-caryophyllene, α-humulene, and α-copaene (100:10:1, β-caryophyllene concentration of 0.17 µg/µl) were loaded on dynamic devices. VOCs were captured every three minutes for 15 min, with 11 replicates. TD-GC-MS analyses showed a steady release of these semiochemicals throughout the recorded time (Fig. 3), thus corroborating earlier findings with other semiochemicals ^13,14^ and indicating these devices are suitable for behavioral experiments.

**Figure 3.**
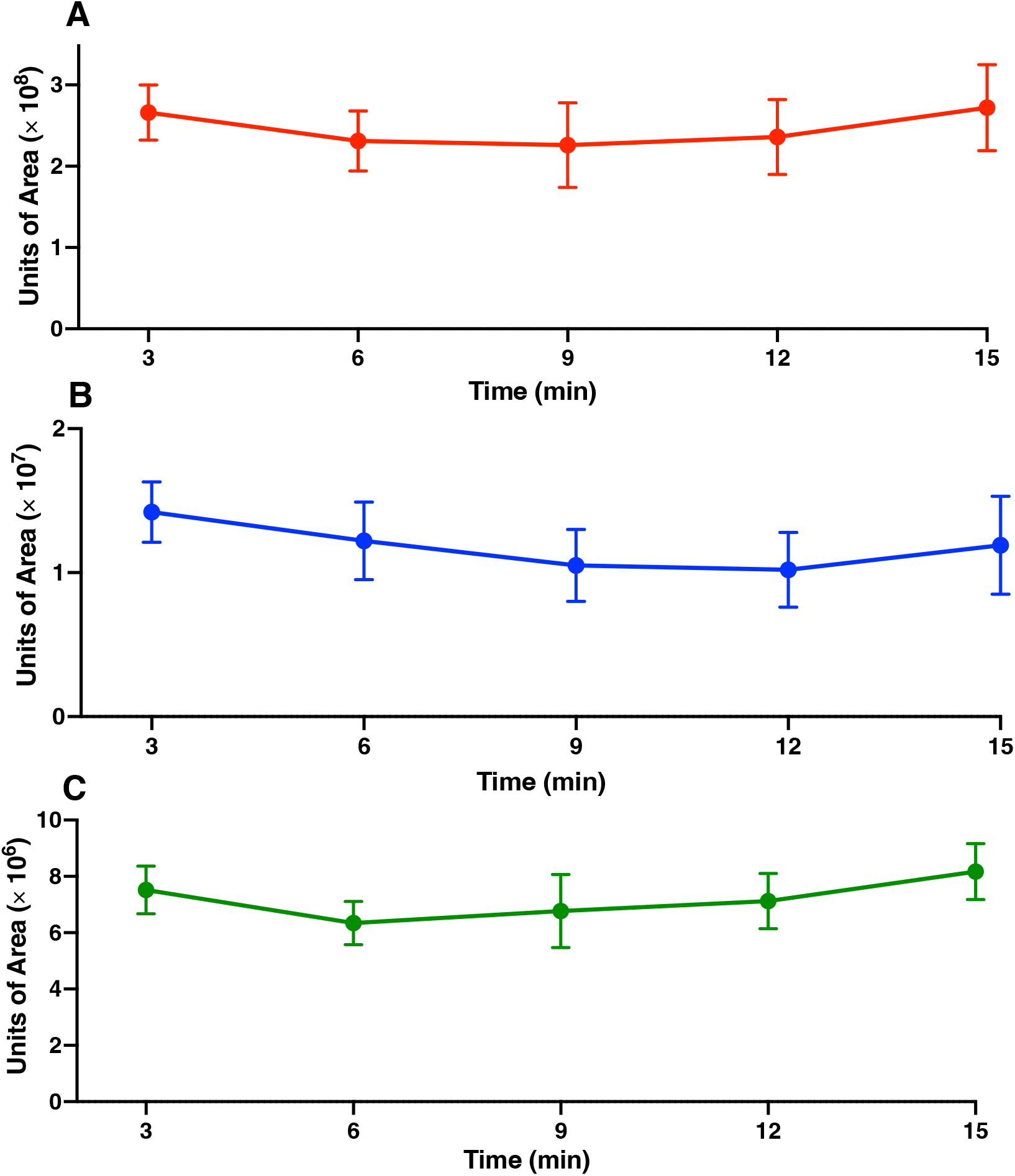
Time-course of sesquiterpene release from dynamic devices. Amounts of (A) β-caryophyllene, (B) α-humulene, and (C) α-copaene released from a dynamic device loaded with a mixture of these three sesquiterpenes in the ratio 100:10:1 based on the dose of β-caryophyllene (0.17 g/µl). Headspace volatiles from the odor chambers were captured every 3 min for 15 min and analyzed by TD-GC-MS. The y-axis’ “units of area” represent the areas generated by the total ion chromatograms for each sesquiterpene. Bars for each data point represent sem.

### Repellence tests of the β-caryophyllene isomers

We surmised whether α-copaene or α-humulene at the rates released from the *A. thaliana* line overexpressing synthase At5g23960 would affect ACP behavioral responses to β-caryophyllene. There was no significant difference in ACP residence times in the odor and control fields when α-humulene was tested at the source dose of 0.017 µg/µl, i.e., 1/10^th^ of β-caryophyllene’s dose (n = 101, p = 0.3145, Wilcoxon test). By contrast, ACP females were significantly repelled by α-copaene at a dose 100x lower than β-caryophyllene in the emanations from the transgenic plant, i.e., 1.7 ng/µl (n = 106, p = 0.0168, Wilcoxon test; mean residence times in the treatment and control fields were 4.25 ± 0.31 min and 5.75 ± 0.31 min, respectively). Intriguingly, it has been reported that 1 µg/µl of α-copaene repels ACP, ^15^ a dose 588x higher than the dose we tested. We surmised this discrepancy could be derived from different methods for releasing α-copaene. However, the release rate reported for the active dose (32 ± 0.2 µg/min) ^15^ was 40,000x higher than the release rate of α-copaene in our experiments (n = 17, 0.80 ± 0.04 ng/min). Dose-response relationship analysis demonstrated that α-copaene is repellent at 0.9 to 2.1 ng/µl (source doses). We tested α-copaene at eight doses from 0.1 to 2.9 ng/µl. Specifically, 0.1 ng/µl (n = 99, p = 0.4262), 0.5 ng/µl (n = 107, p = 0.7954), 0.9 ng/µl (n = 129, p = 0.0042), 1.3 ng/µl (n = 126, p = 0.0052), 1.7 ng/µl (n = 106, p = 0.0168), 2.1 ng/µl (n = 117, p = 0.0210), 2.5 ng/µl (n = 107, p = 0.3124), 2.9 ng/µl (n = 103, p = 0.6154; Wilcoxon tests). At doses ranging from 0.9 to 2.1 ng/µl, ACP females spent significantly more time in the control than in the α-copaene fields (Fig. 4).

**Figure 4.**
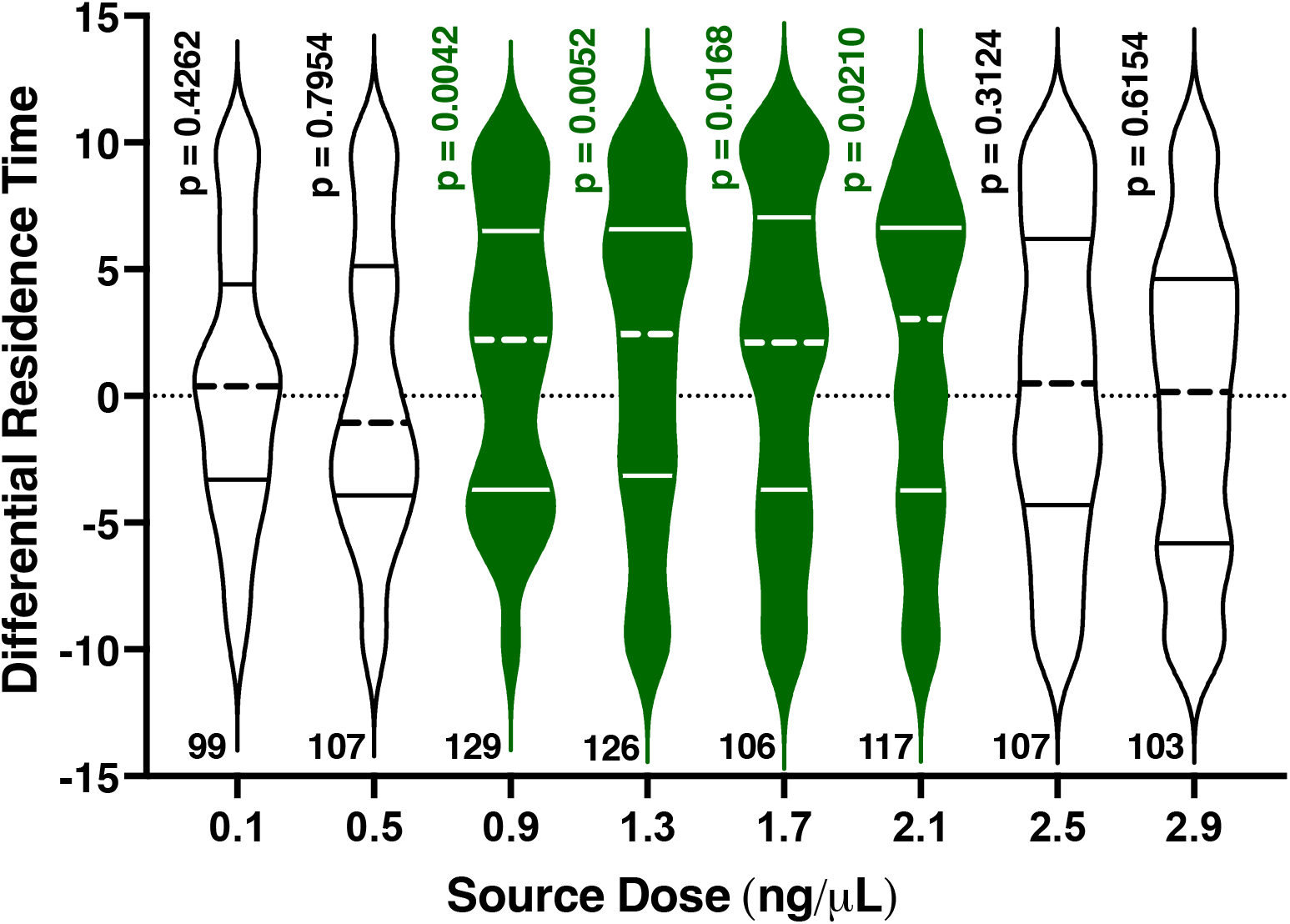
Behavioral responses of ACP females to α-copaene in a multi-choice olfactometer. Violin plots represent the differential residence times, i.e., the residence times in the arena’s control fields minus the residence times in the treatment fields. Statistical significance was determined by comparing the original residence times in the treatment and control fields (Wilcoxon test). Medians and quartiles are denoted by dashed and solid lines, respectively. The violin plots corresponding to significantly different means of residence times (p < 0.05) were colored green. The numbers of replicates are shown below each violin plot.

In conclusion, at the doses calculated based on the ratio of sesquiterpenes released from *A. thaliana* overexpressing the enzyme involved in synthesizing these sesquiterpenes, α-copaene and β-caryophyllene are repellents, but α-humulene is not. Next, we examined whether the active repellents would synergize.

### Evaluating binary and tertiary mixtures

Surprisingly, there was no significant difference in the ACP residence times in the control and treatment fields when a mixture of 100:1 β-caryophyllene (0.17 µg/µl) and α-copaene (1.7 ng/µl) was tested (n = 100, p = 0.6841, Wilcoxon test) (Fig. 5A). ACP females spent 5.15 ± 0.29 min and 4.85 ± 0.29 min in the treatment and control fields, respectively. Because β-caryophyllene and α-copaene are active when tested individually at these doses (Fig. 2A, Fig. 4), we hypothesized that they saturated the olfactory system when combined. To test this hypothesis, we measured the repellence with a mixture of 100:1 β-caryophyllene and α-copaene at the doses of 0.13 µg/µl and 1.3 ng/µl, respectively. ACP females spent significantly more time in the control than in the treatment fields (n = 108, p = 0.0391; Wilcoxon test) (Fig. 5B). ACP females spent 4.51 ± 0.26 min and 5.49 ± 0.26 min in the treatment and control fields, respectively. Because β-caryophyllene per se is not active at this dose (0.13 µg/µl, Fig. 2 B), we concluded that α-copaene plays a more crucial role in the repellence elicited by the *Arabidopsis* transgenic line, which has enormous potential for practical applications.

**Figure 5.**
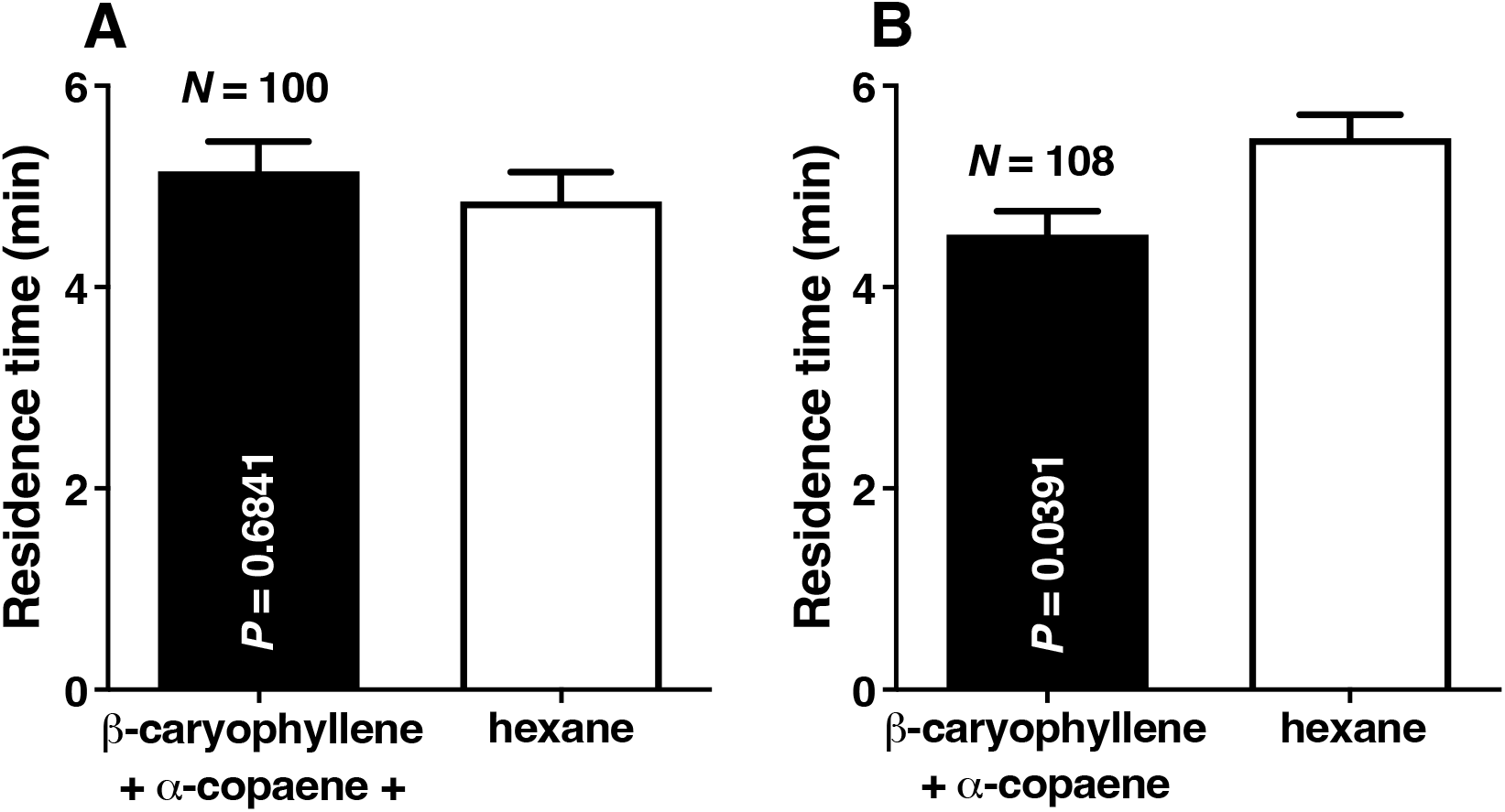
Behavioral responses of ACP females challenged with binary mixtures. (A) ACP behavioral responses to a mixture of β-caryophyllene (0.17 µg/µl) and α-copaene (1.7 ng/µl). β-caryophyllene and α-copaene elicited significant repellence at these doses when tested individually (see above) but not when combined (p = 0.6841). (B) ACP females were significantly repelled (p = 0.0391) by a mixture with lower doses of β-caryophyllene (0.13 µg/µl) and α-copaene (1.3 ng/µl).

Although α-humulene was not active per se at the dose determined by the β-caryophyllene/ α-humulene ratio emitted by the *Arabidopsis* transgenic line (Fig. 1), we tested a tertiary mixture, given the unexpected interaction between α-copaene and β-caryophyllene (Fig. 5A). ACP females spent significantly less time in the field permeated with a 100:10:1 mixture of β-caryophyllene (0.13 µg/µl), α-humulene (0.013 µg/µl), α-copaene (1.3 ng/µl) than in the control fields (n = 112, p = 0.0224, Wilcoxon test).

Whether the two repellents (β-caryophyllene and α-copaene) act on the same olfactory neuron on the ACP sensilla is unknown. Single unit recordings showed that β-caryophyllene elicits the most robust response from tested plant compounds ^16,17^ by activating the sensory neuron B in the antennal rhinarial plate 7 (RP7), ^17^ but α-copaene has never been tested. One may speculate that being isomers, α-copaene and β-caryophyllene would act on the same neuron. However, α-humulene is also a β-caryophyllene isomer and activates the sensory neuron B in RP2. ^17^

We concluded that two of the sesquiterpenes enriched in the *Arabidopsis* transgenic line,^8^β-caryophyllene and α-copaene, are repellents against ACP, but humulene is not. Of note, α-copaene is active at doses ca. 100 times lower than the active doses of β-caryophyllene.

### Copaiba oil repels ACP

*Copaifera* species (Fabaceae), also known in Brazil as “copaiba,” are well-known sources of sesquiterpenes for applications in the pharmaceutical and cosmetic industries. ^18^ Interestingly, β-caryophyllene, α-copaene, and α-humulene appear to be the chemical markers in *Copaifera* volatile oils. ^19^ Although the contents vary remarkably among *Copaifera* species and variants, ^20^ we noticed that the chromatogram profiles of the oil derived from a species native to Brazil, *Copaifera langsdorffii*, ^18^ resemble that of the *Arabidopsis* transgenic line (Fig. 1). GC-MS analyses showed that a sample of refined copaiba oil from Beraca contained β-caryophyllene (517.09 ± 9.54 mg/ml), α-humulene (60.33 ± 0.71 mg/ml), and α-copaene (28.26 ± 0.50 mg/ml). A shoulder peak after the β-caryophyllene peak was tentatively identified as α-bergamotene (CAS# 13474-59-4).

To test the repellence activity of the copaiba oil, we diluted the commercial product 3,500X to generate a sample with approximately the same active dose of β-caryophyllene (Fig. 2A). The diluted copaiba oil repelled ACP females, which spent significantly more time in the control (6.02 ± 0.28 min) than in the treatment fields (3.98 ± 0.28 min, n = 115, p = 0.0005, Wilcoxon test). The ACP responses to control and treatments did not significantly differ when the copaiba oil was diluted 4,000x or 3,000x (n = 122, p = 0.1402; n = 99, p = 0.3939, respectively).

Next, we measured the emission rates of the active repellents β-caryophyllene and α-copaene by TD-GC-MS (Fig. 6). The emission rates of these repellents captured in the headspace generated by 3,500x diluted samples (n = 8) were β-caryophyllene (67 ± 3 ng/min) and α-copaene (8.4 ± 0.1 ng/min). The ratio of α-copaene/β-caryophyllene was 0.128 ± 0.006 (n = 8).

**Figure 6.**
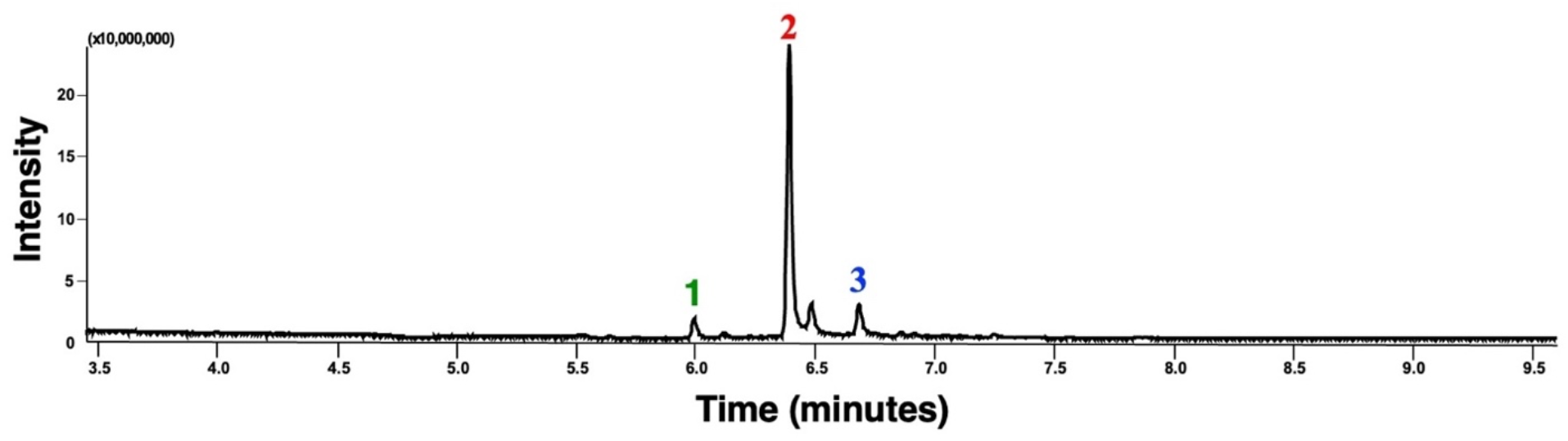
Chromatogram obtained with a sample of refined copaiba oil. This TD-GC-MS chromatogram was obtained with a commercial copaiba oil (Beraca) sample. The most prominent peaks were identified as (1) α-copaene, (2) β-caryophyllene, and (3) α-humulene based on their mass spectra, retention times, and comparison with authentic standards. This profile resembles the chromatogram obtained with VOCs from the *Arabidopsis* transgenic line (Fig. 1).

### Concluding Remarks

As expected, the *Arabidopsis* transgenic line ^8^ that overexpresses the sesquiterpene synthase At5g44630 ^10^ yields β-caryophyllene, α-humulene, α-copaene, and traces amounts of β-elemene. With TD-GC-MS, we demonstrated that β-caryophyllene, α-humulene, and α-copaene are released in an approximate ratio of 100:10:1. Our studies further corroborate that β-caryophyllene released from a dynamic device loaded with 0.17 µg/µl, a dose equivalent to 1 µg/µl in a static device (open glass vial), ^8^ repels ACP females. The dynamic device loaded with 1,000 µl of 0.17 µg/µl releases β-caryophyllene at 0.044 ± 0.013 µg/min. Interestingly, α-copaene is an even more potent repellent, active at 0.80 ± 0.04 ng/min, i.e., a 55x lower amount. Additionally, α-copaene showed a more comprehensive range of active concentrations (0.9 to 2.1 ng/µl) than β-caryophyllene. Previously, α-copaene has been reported as a repellent against ACP, ^15^ but intriguingly, the tested dose (1µg/µl) was more than 500x higher than the active range of concentrations we found in these studies. Although we did not find evidence of synergistic activity between α-copaene and β-caryophyllene, higher doses of the two repellents combined may overload the olfactory system. It is worth noting that upon release into the environment (whether from plants or chemical devices), β-caryophyllene and α-copaene undergo degradation, with β-caryophyllene depleting more rapidly, ^21^ partly because of the two double bonds (as opposed to a single double bond in α-copaene, Fig. 1). It has been estimated that the rate constant for β-caryophyllene reaction with ozone is ca. 38x the rate constant for α-copaene. ^21^ Therefore, α-copaene seems more suitable for possible repellent chemical formulations. The data presented here show that copaiba oil is an economically viable source of these sesquiterpene repellents, which may be explored to develop slow-releasing devices to repel ACP.

Our studies further demonstrate that transgenic plants overexpressing β-caryophyllene and α-copaene ^8,9^ may become invaluable tools in ACP management strategies. For example, combined with plants attractive to ACP (e.g., *M. paniculata*, ^6^ *Bergera koenigii* ^*5*^), these transgenic plants may be applied in push-pull strategies. ^4^

## Materials and Methods

### Insects

The adults of *D. citri* used in bioassays were derived from a *Ca*. Liberibacter spp-free colony maintained for several generations in a greenhouse at FUNDECITRUS. The insects were kept in metal cages covered with anti-aphid mesh (200 mesh). Seedlings of *M. paniculata* were pruned to a height of 30 cm to stimulate the emergence of shoots. After sprouting (0.5 to 1.0 cm in length), the seedlings were placed in cages, into which 400 mated adults were released 10–15 days after emergence. The adults were kept in cages for 7 days for oviposition on the seedlings. After that, the adults were removed, and the seedlings containing eggs were retained in the cages for nymphal development. Cages containing fourth-and fifth-instar nymphs were transported to an air-conditioned room (temperature 25 ± 2°C, relative humidity 60% ± 10%, and photoperiod 14-h L:10-h D) for the emergence of adults and maintenance of insects under environmental conditions similar to those used in behavioral assays. Newly emerged adults were removed from cages daily and confined on *M. paniculata* seedlings coated with a tulle-like material for mating and age control of the insects. We used mated females between 7 and 15 days after emergence for all bioassays.

### Behavioral measurements and statistical analysis

All repellence responses were measured with a previously described multi-choice olfactometer ^22^ operated in a climate-controlled room at 25 ± 2 °C, 65 ± 10% relative humidity, and 3,000 lux luminosity. In brief, we used an acrylic 4-arm olfactometer (30.0 × 30.0 × 2.5 cm; length × width × height, respectively) with a transparent acrylic lid and a yellow laserjet print paper (lightness, 84.8; chroma, 98.7; and hue angle, 95.7) under the device ^22^. Compressed air (charcoal filtered and subsequently humidified) was connected to a stainless-steel line. It split into four individual 0.635-cm-diameter polytetrafluoroethylene (PTFE) tubes (Sigma-Aldrich, Bellefonte, PA, USA) connected to four flowmeters [0.1–1 liter per minute (LPM), Brooks Instruments, Hatfield, PA, USA], adjusted to an airflow of 0.1 LPM. Each PTFE tube was connected to one horizontal glass chamber (20 cm length × 6 cm internal diameter) containing an odor source, and each airflow converged through PTFE tubes to one of the four device arms. Data recorded for the time of residence of individual insects in each odor field (treatment or control) were first analyzed to determine normality using the Shapiro-Wilk test with the Prism 10.1.1 software (GraphPad, La Jolla, California, USA). Data that did not meet the assumption of normal distribution were analyzed using the non-parametric Wilcoxon matched-pairs signed rank test. Descriptive statistics are provided in the figure and figure legends. Means are presented along the standard error of the means (mean ± sem).

### Semiochemicals releasing devices

Previously, we have used static devices to release semiochemicals when measuring their repellence activity ^8^. Specifically, the test compounds like β-caryophyllene^8^ were released from open glass vials (2 ml) containing hexane solutions (100 µl) of the semiochemicals. To test mixtures, it is critical to release semiochemicals in the desired ratio despite differences in the volatility of the mixture’s constituents. We used a slow-release (“dynamic”) device inspired on the “wick-baits” ^14^ designed to release a steady flow of semiochemical mixtures faithfully representing the proportion of the components in the mixture. ^13^ The “dynamic” device was comprised of a 2 ml screw-top clear glass vial (#27329, Supelco, Bellefonte, PA, USA) capped with a blue polypropylene screw cap (#SU860092, Supelco). A 4 cm PTFE tubing (ID 1.58 mm; OD 2.1 mm, Supelco, #20531) filled with cotton yarn wick (“barbante cru 100% algodão,” São Francisco Industrial & Comercial Textil Ltd., Piracicaba, São Paulo, Brazil) was inserted through the vial’s cap PTFE septum and placed 1 mm from the bottom of the vial. The cotton yarn wick was prepared by washing for 5 min in hexane, 3 times and then dried up before inserting into the tubing. The vials were loaded with 1 ml of hexane solutions of individual compounds or mixtures.

### Chemical analysis

For volatile collections, each outlet of the glass odor chambers was connected to a homemade two-way glass valve with PTFE connectors. Each side of the valve was connected to a Tenax^®^ TA tube for capturing volatile organic compounds (VOCs). The outlets of the two Tenax tubes were connected to a second two-way glass valve, and the final outlet tube was connected to a vacuum line. These two-way valve systems allowed continuous sampling for time course analysis without disrupting the headspace or the airflow. After a specific time (e.g., 3 min), the airflow was diverted to the oppositive side to trap VOCs for the next time measurement (e.g., 3-6 min). The Tenax tube was replaced with a clean one for the subsequent collection (e.g., 6-9 min); the trapped VOCs were analyzed as described below. The specific collection for each experiment is described in the figure legends. Multiple glass odor chambers (N = 7-11) were used for replicates.

Volatile organic compounds (VOCs) captured in headspace samples were analyzed using thermal desorber–gas chromatography-mass spectrometry (TD-GC-MS) equipment. The volatile compounds were thermally desorbed from Tenax^®^ TA (0.635 × 8.89 cm glass tubes containing 200 mg of 2.6-diphenyl-*p*-phenylene oxide 35-60 mesh; Sigma-Aldrich, Bellefonte, PA, USA) in the TD equipment (ULTRA-xr Thermal Desorber, Markes International Ltd., Llantrisant, UK) under a 50 mL/min helium gas flow heated to 280°C for 5 min. The VOCs were directed to Tenax^®^ TA-based cold traps and were cryogenically captured at a temperature of -20°C. Subsequently, the cold traps were desorbed at 300°C for 3 min — the transfer lines from the cold traps to the gas chromatographs were set at 200°C. For GC-MS analyses, the following instruments were used: Shimadzu’s QP2010-plus (Shimadzu Corporation, Kyoto, Japan) and Thermo’s TRACE 1310/TSQ 9000 (Thermo Fisher Scientific (Austin, TX, USA). The Shimadzu’s GC instrument was equipped with Rxi-5ms columns (10 m, 0.10 mm, 0.10 μm film thickness or 30 m, 0.25 mm, 0.25 μm film thickness; Restek, Bellefonte, PA). The Thermo’s GC (TRACE 1310) had a 10 m Rxi-5ms column. The Shimadzu’s and Thermo’s GC/MS transfer lines were kept at 250°C. The ovens of the GCs equipped with 10 m columns were operated at the initial temperature of 40°C and a rate of 20°C/min until 250°C, and then it was held at the final temperature for 2 min [40-20-250(2)]. The temperature program for GCs with 30 m columns was 40-7-250(5). Mass spectra were obtained at a collision energy of 70 eV, and all analyses were recorded in total ion chromatogram (TIC) mode, with a scanning range of *m/z* 40 to 450. Authentic compounds for chemical identification and quantification (calibration curves) were injected into a Calibration Solution Loading Ring (CSLR, Tenax^®^ TA) at 0.1 ml/min for 5 min. The authentic compounds loaded on Tenax^®^ TA tubes were desorbed, cryogenically captured, and subsequently injected into GCs following the sampling protocol described above for natural products. Copaiba oil was initially analyzed by injecting 0.5 µl hexane samples into GC-MS for chemical identification and estimation of the ratios. The oven was operated at 60-15-300(2), helium (0.4 ml/min) was used as the carrier gas, and the injector in the split mode (20:1) was set at 250 °C. The emission rates of copaiba oil and other individual repellents were measured with calibration curves obtained with authentic standards using the thermal desorber-gas chromatography-mass spectrometry technique.

### Chemicals

β-Caryophyllene (≥ 98.5% purity) and α-humulene (≥ 96.0% purity) were acquired from Sigma Aldrich (Steinheim, Germany). α-Copaene (94% purity) and hexane (≥ 99.0% purity) were acquired from TRC (Toronto, Canada) and Merck (Darmstadt, Germany), respectively. Refined copaiba oil was purchased from BERACA Ingredientes Naturais (BR03310BX45, batch 09183310R, Levilandia-Anamindeuz, Para, Brazil).

### Terpene synthase overexpressing line of *Arabidopsis thaliana*

Seeds of the previously reported *A. thaliana* line^8^ overexpressing the β-caryophyllene synthase At5g23960 ^10^ were grown to the flouring state on soil in a controlled climate chamber at 22 °C and 55% relative humidity for up to seven weeks under a photoperiod 14-h L:10-h D, and 3,000 lux.

## Acknowledgments

We thank Victoria Esperança, Giulia Lopes de Souza, and Isabela Cristina da Silva Teixeira (FUNDECITRUS) for their assistance with chemical and behavioral analysis, Drs. Edson Rodrigues Filho (LaBioMMi, Department of Chemistry, FFCLRP, UFSCar, São Carlos) and Luiz A. B. de Moraes (Department of Chemistry, FFCLRP, USP, Ribeirão Preto, Brazil) for sharing GC-MS equipment used in the initial analysis, and Dr. Marcelo P. Miranda (FUNDECITRUS) for discussion. This work was partially supported by the São Paulo State Research Foundation (FAPESP, 2015/07011-3 and 2017/21460-0) and the Fund for Citrus Protection (FUNDECITRUS) under a research agreement with the University of California, Davis (#201600147).

## Author contributrions

W.S.L. R.F.M. and H.X.L.V. designed research and analyzed data. R.F.M., H.X.L.V., R.A.G.L., T.A.M., T.T.A., M.C.-S., N.A.W., and L.P. performed research. J.K.B. and V.P.R. provided reagents. W.S.L. wrote the manuscript. All authors reviewed and approved the final version of the manuscript.

## Additional Information

A dataset with raw data used to generate figures accompanies this paper at xxxx

## Data Availability

The manuscript and supporting files include all data generated during this study.

## References

1 Bove, J. M. Huanglongbing: A destructive, newly-emerging, century-old disease of citrus. J Plant Pathol 88, 7–37 (2006).

2 Coletta-Filho, H. D. et al. First Report of the Causal Agent of Huanglongbing (“Candidatus Liberibacter asiaticus”) in Brazil. Plant Dis 88, 1382, doi:10.1094/PDIS.2004.88.12.1382C (2004).

3 FUNDECITRUS. Levantamento da incidência das doenças dos citros: greening, CVC e cancro cítrico, < https://www.fundecitrus.com.br/pdf/levantamentos/Relatorio_Levantamento_de_doencas_2024_completo.pdf > (2024).

4 Cook, S. M., Khan, Z. R. & Pickett, J. A. The use of push-pull strategies in integrated pest management. Annu Rev Entomol 52, 375–400, doi:10.1146/annurev.ento.52.110405.091407 (2007).

5 Eduardo, W. I. et al. Push-pull and kill strategy for Diaphorina citri control in citrus orchards. Entmol. Exp. Appl. 171, 287–299, doi:10.1111/eea.13273 (2023).

6 Tomaseto, A. F. et al. Orange jasmine as a trap crop to control Diaphorina citri. Sci Rep 9, 2070, doi:10.1038/s41598-019-38597-5 (2019).

7 Cifuentes-Arenas, J. C., Beattie, G. A. C., Peña, L. & Lopes, S. A. Murraya paniculata and Swinglea glutinosa as Short-Term Transient Hosts of ‘Candidatus Liberibacter asiaticus’ and Implications for the Spread of Huanglongbing. Phytopathology® 109, 2064–2073, doi:10.1094/phyto-06-19-0216-r (2019).

8 Alquezar, B. et al. beta-caryophyllene emitted from a transgenic Arabidopsis or chemical dispenser repels Diaphorina citri, vector of Candidatus Liberibacters. Sci Rep 7, 5639, doi:10.1038/s41598-017-06119-w (2017).

9 Alquezar, B. et al. Engineered Orange Ectopically Expressing the Arabidopsis beta-Caryophyllene Synthase Is Not Attractive to Diaphorina citri, the Vector of the Bacterial Pathogen Associated to Huanglongbing. Front Plant Sci 12, 641457, doi:10.3389/fpls.2021.641457 (2021).

10 Tholl, D., Chen, F., Petri, J., Gershenzon, J. & Pichersky, E. Two sesquiterpene synthases are responsible for the complex mixture of sesquiterpenes emitted from Arabidopsis flowers. Plant J 42, 757–771, doi:10.1111/j.1365-313X.2005.02417.x (2005).

11 Zaka, S. M., Zeng, X.-N., Holford, P. & Beattie, G. A. C. Repellent effect of guava leaf volatiles on settlement of adults of citrus psylla, Diaphorina citri Kuwayama, on citrus. Insect Science 17, 39–45, doi:10.1111/j.1744-7917.2009.01271.x (2010).

12 Chen, F. et al. Biosynthesis and emission of terpenoid volatiles from Arabidopsis flowers. Plant Cell 15, 481–494, doi:10.1105/tpc.007989 (2003).

13 Birgersson, G. in Annual Meeting of the International Society of Chemical Ecology 56 (Stockholm, Sweeden, 2015).

14 Birgersson, G., Dalusky, M. J., Espelie, K. E. & Berisford, C. W. Pheromone Production, Attraction, and Interspecific Inhibition among Four Species of Ips Bark Beetles in the Southeastern USA. Psyche: A Journal of Entomology 2012, 532652, doi:10.1155/2012/532652 (2012).

15 Pancarte, C., Altamimi, R., Fall, M. N. D. & Martini, X. Repellency of volatiles from Martinique island guava varieties against Asian citrus psyllids. Arthropod-Plant Interactions 16, 341–348, doi:10.1007/s11829-022-09901-4 (2022).

16 Coutinho-Abreu, I. V., Forster, L., Guda, T. & Ray, A. Odorants for surveillance and control of the Asian Citrus Psyllid (Diaphorina citri). PLoS One 9, e109236, doi:10.1371/journal.pone.0109236 (2014).

17 Coutinho-Abreu, I. V., McInally, S., Forster, L., Luck, R. & Ray, A. Odor coding in a disease-transmitting herbivorous insect, the Asian citrus psyllid. Chem Senses 39, 539–549, doi:10.1093/chemse/bju023 (2014).

18 Sousa, J. P. et al. Validation of a gas chromatographic method to quantify sesquiterpenes in copaiba oils. J Pharm Biomed Anal 54, 653–659, doi:10.1016/j.jpba.2010.10.006 (2011).

19 Pinto, A. C. et al. Separation of Acid Diterpenes of Copaifera cearensis Huber ex Ducke by Flash Chromatography Using Potassium Hydroxide Impregnated Silica Gel. Journal of the Brazilian Chemical Society, doi:10.1590/S0103-50532000000400005 (2000).

20 Veiga Junior, V. F., Rosas, E. C., Carvalho, M. V., Henriques, M. G. & Pinto, A. C. Chemical composition and anti-inflammatory activity of copaiba oils from Copaifera cearensis Huber ex Ducke, Copaifera reticulata Ducke and Copaifera multijuga Hayne--a comparative study. J Ethnopharmacol 112, 248–254, doi:10.1016/j.jep.2007.03.005 (2007).

21 Pollmann, J., Ortega, J. & Helmig, D. Analysis of Atmospheric Sesquiterpenes: Sampling Losses and Mitigation of Ozone Interferences. Environmental Science & Technology 39, 9620–9629, doi:10.1021/es050440w (2005).

22 Volpe, H. X. L. et al. Behavioral responses of Diaphorina citri to host plant volatiles in multiple-choice olfactometers are affected in interpretable ways by effects of background colors and airflows. PLoS One 15, e0235630, doi:10.1371/journal.pone.0235630 (2020).

