## Additional Information for "α-Copaene is a potent repellent against the Asian Citrus Psyllid *Diaphorina citri*"

### Dataset with the raw data used to generate figures 2, 3, 4, and 5

| Figure 2A | | Figure 2B | | Figure 2C | |
|---|---|---|---|---|---|
| **Caryophyllene** | **Control** | **Caryophyllene** | **Control** | **Caryophyllene** | **Control** |
| 0 | 10 | 0.15 | 9.85 | 3.583333 | 6.416667 |
| 3.166667 | 6.833333 | 0.55 | 9.45 | 4.866667 | 5.133333 |
| 3.133333 | 6.866667 | 3.05 | 6.95 | 0 | 10 |
| 10 | 0 | 0 | 10 | 7.083333 | 2.916667 |
| 10 | 0 | 6.83 | 3.17 | 6.55 | 3.45 |
| 7.633333 | 2.366667 | 1.62 | 8.38 | 0.95 | 9.05 |
| 0 | 10 | 5.78 | 4.22 | 1.683333 | 8.316667 |
| 6.25 | 3.75 | 5.67 | 4.33 | 7.15 | 2.85 |
| 0 | 10 | 2.75 | 7.25 | 5.466667 | 4.533333 |
| 6.233333 | 3.766667 | 1.07 | 8.93 | 4.316667 | 5.683333 |
| 0 | 10 | 6.58 | 3.42 | 4.966667 | 5.033333 |
| 10 | 0 | 8.18 | 1.82 | 5.233333 | 4.766667 |
| 0 | 10 | 2.02 | 7.98 | 10 | 0 |
| 10 | 0 | 6.48 | 3.52 | 5.183333 | 4.816667 |
| 6.933333 | 3.066667 | 0 | 10 | 6.25 | 3.75 |
| 0.25 | 9.75 | 2.6 | 7.4 | 0.883333 | 9.116667 |
| 0 | 10 | 0.42 | 9.58 | 1.233333 | 8.766667 |
| 10 | 0 | 6.6 | 3.4 | 10 | 0 |
| 0 | 10 | 5.25 | 4.75 | 0.716667 | 9.283333 |
| 0 | 10 | 4.75 | 5.25 | 0.95 | 9.05 |
| 6.066667 | 3.933333 | 10 | 0 | 4.583333 | 5.416667 |
| 4.533333 | 5.466667 | 6.93 | 3.07 | 0.85 | 9.15 |
| 4.566667 | 5.433333 | 6.28 | 3.72 | 0.366667 | 9.633333 |
| 0 | 10 | 2.15 | 7.85 | 5.9 | 4.1 |
| 0 | 10 | 4.57 | 5.43 | 2.783333 | 7.216667 |
| 0 | 10 | 4.03 | 5.97 | 0 | 10 |
| 4.516667 | 5.483333 | 0 | 10 | 0 | 10 |
| 10 | 0 | 6.03 | 3.97 | 6.033333 | 3.966667 |
| 10 | 0 | 10 | 0 | 8.966667 | 1.033333 |
| 3.783333 | 6.216667 | 2.8 | 7.2 | 6.5 | 3.5 |
| 0 | 10 | 4.28 | 5.72 | 6.6 | 3.4 |

| | | | | | |
|---|---|---|---|---|---|
| 0 | 10 | 5.77 | 4.23 | 0.966667 | 9.033333 |
| 0.633333 | 9.366667 | 0 | 10 | 0 | 10 |
| 9.8 | 0.2 | 0 | 10 | 9.516667 | 0.483333 |
| 4.066667 | 5.933333 | 9.47 | 0.53 | 10 | 0 |
| 0 | 10 | 8.67 | 1.33 | 6.866667 | 3.133333 |
| 0 | 10 | 1.67 | 8.33 | 10 | 0 |
| 0.55 | 9.45 | 8.72 | 1.28 | 7.95 | 2.05 |
| 5.25 | 4.75 | 4.03 | 5.97 | 4.433333 | 5.566667 |
| 0 | 10 | 9.85 | 0.15 | 8.85 | 1.15 |
| 9.533333 | 0.466667 | 0.25 | 9.75 | 6.233333 | 3.766667 |
| 1.333333 | 8.666667 | 1.18 | 8.82 | 5.833333 | 4.166667 |
| 10 | 0 | 6.67 | 3.33 | 7.116667 | 2.883333 |
| 2.05 | 7.95 | 6.67 | 3.33 | 4.1 | 5.9 |
| 4.166667 | 5.833333 | 3.43 | 6.57 | 0.2 | 9.8 |
| 9.633333 | 0.366667 | 0 | 10 | 1.883333 | 8.116667 |
| 10 | 0 | 0 | 10 | 5.166667 | 4.833333 |
| 2.6 | 7.4 | 2.53 | 7.47 | 1.983333 | 8.016667 |
| 3.933333 | 6.066667 | 8.7 | 1.3 | 5.833333 | 4.166667 |
| 6.633333 | 3.366667 | 6.83 | 3.17 | 6.2 | 3.8 |
| 9.766667 | 0.233333 | 0.32 | 9.68 | 6.283333 | 3.716667 |
| 0 | 10 | 0.57 | 9.43 | 2.15 | 7.85 |
| 0 | 10 | 0.78 | 9.22 | 1.666667 | 8.333333 |
| 5.716667 | 4.283333 | 0.38 | 9.62 | 0 | 10 |
| 1.683333 | 8.316667 | 10 | 0 | 1.15 | 8.85 |
| 7.466667 | 2.533333 | 1.03 | 8.97 | 0.95 | 9.05 |
| 2.55 | 7.45 | 5.87 | 4.13 | 1.2 | 8.8 |
| 0 | 10 | 7.9 | 2.1 | 2.35 | 7.65 |
| 7.033333 | 2.966667 | 10 | 0 | 8.416667 | 1.583333 |
| 2.3 | 7.7 | 1.23 | 8.77 | 3 | 7 |
| 0 | 10 | 10 | 0 | 6.05 | 3.95 |
| 4.916667 | 5.083333 | 1.42 | 8.58 | 6.816667 | 3.183333 |
| 9.5 | 0.5 | 5.6 | 4.4 | 0 | 10 |
| 0 | 10 | 4.85 | 5.15 | 0 | 10 |

| | | | | | |
|---|---|---|---|---|---|
| 5.65 | 4.35 | 10 | 0 | 1.116667 | 8.883333 |
| 0 | 10 | 2.68 | 7.32 | 6.133333 | 3.866667 |
| 0.316667 | 9.683333 | 0.22 | 9.78 | 5.2 | 4.8 |
| 1.25 | 8.75 | 5.25 | 4.75 | 10 | 0 |
| 0.483333 | 9.516667 | 3.15 | 6.85 | 2.9 | 7.1 |
| 2.566667 | 7.433333 | 2.6 | 7.4 | 8.45 | 1.55 |
| 9.766667 | 0.233333 | 0.6 | 9.4 | 7.65 | 2.35 |
| 9.1 | 0.9 | 8.57 | 1.43 | 8.3 | 1.7 |
| 3.15 | 6.85 | 2.87 | 7.13 | 8.166667 | 1.833333 |
| 4.95 | 5.05 | 2.47 | 7.53 | 4.366667 | 5.633333 |
| 4.55 | 5.45 | 8.98 | 1.02 | 7.333333 | 2.666667 |
| 0 | 10 | 5 | 5 | 1.483333 | 8.516667 |
| 0 | 10 | 6.63 | 3.37 | 1.016667 | 8.983333 |
| 0.433333 | 9.566667 | 0 | 10 | 3.433333 | 6.566667 |
| 0 | 10 | 4.12 | 5.88 | 1.15 | 8.85 |
| 0.666667 | 9.333333 | 4.95 | 5.05 | 0.733333 | 9.266667 |
| 10 | 0 | 10 | 0 | 8.933333 | 1.066667 |
| 0.616667 | 9.383333 | 0 | 10 | 6.583333 | 3.416667 |
| 6.716667 | 3.283333 | 2.67 | 7.33 | 6.5 | 3.5 |
| 0 | 10 | 2.3 | 7.7 | 10 | 0 |
| 1.3 | 8.7 | 3.25 | 6.75 | 0 | 10 |
| 7.883333 | 2.116667 | 9.1 | 0.9 | 0.383333 | 9.616667 |
| 0.883333 | 9.116667 | 0 | 10 | 0 | 10 |
| 9.6 | 0.4 | 10 | 0 | 0 | 10 |
| 10 | 0 | 2.67 | 7.33 | 7.266667 | 2.733333 |
| 8.383333 | 1.616667 | 10 | 0 | 2.583333 | 7.416667 |
| 8.783333 | 1.216667 | 6.27 | 3.73 | 4.583333 | 5.416667 |
| 0 | 10 | 0 | 10 | 6.816667 | 3.183333 |
| 5.783333 | 4.216667 | 8.67 | 1.33 | 10 | 0 |
| 1.333333 | 8.666667 | 0 | 10 | 0 | 10 |
| 5.583333 | 4.416667 | 9.2 | 0.8 | 8.066667 | 1.933333 |
| 4.15 | 5.85 | 7.05 | 2.95 | 7.383333 | 2.616667 |
| 10 | 0 | 10 | 0 | 10 | 0 |

| | | | | | |
|---|---|---|---|---|---|
| 8.666667 | 1.333333 | 10 | 0 | 6.366667 | 3.633333 |
| 1.166667 | 8.833333 | 6.43 | 3.57 | 2.683333 | 7.316667 |
| | | 10 | 0 | 0 | 10 |
| | | 10 | 0 | 7.883333 | 2.116667 |
| | | 10 | 0 | 10 | 0 |
| | | 4.52 | 5.48 | 5.9 | 4.1 |
| | | | | 8.1 | 1.9 |
| | | | | 7.216667 | 2.783333 |
| | | | | 0 | 10 |
| | | | | 1.3 | 8.7 |
| | | | | 2.85 | 7.15 |
| | | | | 5.133333 | 4.866667 |
| | | | | 5.783333 | 4.216667 |
| | | | | 2.45 | 7.55 |

| Figure 3A | | | | |
|---|---|---|---|---|
| **Time (min)** | **Caryophyllene** | | | |
| | **Mean (xE8)** | **SEM** | | |
| 3 | 2.66 | 0.34 | | |
| 6 | 2.31 | 0.37 | | |
| 9 | 2.26 | 0.52 | | |
| 12 | 2.36 | 0.46 | | |
| 15 | 2.72 | 0.53 | | |
| **RAW DATA-Caryophyllene** | | | | |
| 3 min | 6 min | 9 min | 12 min | 15 min |
| 8530447 | 11053310 | 15583612 | 8206448 | 9369241 |
| 5781608 | 7167303 | 4845425 | 5597182 | 6078842 |
| 10262541 | 2322740 | 4137058 | 8278420 | 10993271 |
| 9591658 | 4457877 | 8627768 | 11873616 | 11969859 |
| 6870316 | 6288357 | 9319968 | 6620649 | 7386032 |
| 7468203 | 6974767 | 6143931 | 4582375 | 5068477 |
| 4812725 | 9277120 | 4200200 | 4651644 | 6350825 |
| 10990160 | 6770081 | 4729570 | | |
| 3399491 | 5078879 | 3305359 | | |
| | 7418950 | | | |
| | 2932805 | | | |
| **Figure 3B** | | | | |
| **Time (min)** | **Humulene** | | | |
| **Mean (xE7)** | **SEM** | | | |
| 3 | 1.42 | 0.21 | | |
| 6 | 1.22 | 0.27 | | |
| 9 | 1.05 | 0.25 | | |
| 12 | 1.02 | 0.26 | | |
| 15 | 1.19 | 0.34 | | |
| **RAW DATA-Humulene** | | | | |
| 3 min | 6 min | 9 min | 12 min | 15 min |
| 305500185 | 403258211 | 574794931 | 224814575 | 256340259 |
| 197540726 | 445132247 | 123437089 | 183322059 | 212476080 |
| 407160640 | 94899358 | 146400646 | 217051713 | 308990266 |
| 301221722 | 131890232 | 230289298 | 502332751 | 565496076 |
| 256460085 | 218140452 | 377060736 | 234501654 | 265423365 |
| 233645645 | 187920970 | 207492286 | 132794916 | 148932586 |
| 143122708 | 344428638 | 115897680 | 159610252 | 149580804 |
| 414076762 | 214065801 | 177780460 | | |
| 137938013 | 142385338 | 80677668 | | |
| | 282708366 | | | |
| | 74775783 | | | |

| | | | | |
|---|---|---|---|---|
| **Figure 3C** | | | | |
| **Time (min)** | **Copaene** | | | |
| **Mean (xE6)** | **SEM** | | | |
| 3 | 7.52 | 0.85 | | |
| 6 | 6.34 | 0.77 | | |
| 9 | 6.77 | 1.3 | | |
| 12 | 7.12 | 0.98 | | |
| 15 | 8.17 | 0.99 | | |
| **RAW DATA-Copaene** | | | | |
| 3 min | 6 min | 9 min | 12 min | 15 min |
| 15949721 | 19447413 | 27621691 | 8187679 | 9037903 |
| 9613980 | 33978572 | 5338108 | 9048133 | 8695074 |
| 24259965 | 4953987 | 7720424 | 7814254 | 12801520 |
| 14225953 | 5213910 | 8823094 | 25891561 | 31535093 |
| 13935996 | 10734379 | 17946672 | 8252990 | 9876338 |
| 10278038 | 7231610 | 7964407 | 5313738 | 6100143 |
| 8206612 | 17233237 | 5740921 | 7023177 | 5466575 |
| 23833860 | 9190591 | 10160059 | | |
| 7150897 | 7169745 | 3398341 | | |
| | 14546661 | | | |
| | 4221148 | | | |

**Figure 4**

| Differential Residence Times | | | | | | | |
|---|---|---|---|---|---|---|---|
| **0.1 ng/ul** | **0.5 ng/ul** | **0.9 ng/ul** | **1.3 ng/ul** | **1.7 ng/ul** | **2.1 ng/ul** | **2.5 ng/ul** | **2.9 ng/ul** |
| 7.22 | 2.06 | -0.76 | 6.06 | 5.02 | -4.4 | -6.2 | 6.16 |
| 7.86 | 0.36 | -3.42 | -2.46 | -3.1 | -3.74 | 3.8 | 10 |
| 1.46 | -3.72 | -1.68 | -2.54 | -3.58 | -6.88 | 10 | 3 |
| 2.18 | -3.9 | -2.8 | -1.86 | -10 | 1.22 | 2.58 | -5.26 |
| 5.9 | -2.76 | 5.4 | 5.5 | -10 | -10 | -8.4 | -5.82 |
| -0.1 | -1.42 | 10 | 6.5 | 1.5 | 0.02 | -7.82 | 10 |
| 6.26 | -0.22 | -4.04 | 7.22 | 5.42 | 7.66 | 2.02 | 3.88 |
| 0.88 | -5.1 | 10 | 9.26 | 3.76 | -6.12 | -1 | -8.46 |
| -1.66 | 2.9 | -0.1 | 5.56 | 8.64 | -10 | 7.96 | -7.1 |
| 8.64 | 3.52 | 2.28 | -9 | -3.42 | -10 | 5.66 | -5.36 |
| -0.84 | -3.32 | 2.36 | -10 | 10 | 5.8 | -6.02 | 4.66 |
| 1.26 | -2.04 | -2.84 | -7.4 | 10 | 0.16 | 0.36 | 9.88 |
| -5.3 | -10 | 0.96 | -7.52 | -4.06 | 6.8 | -6.74 | -5.94 |
| -0.36 | 7.54 | 7.58 | 6.4 | -1.66 | 6.14 | 8.14 | 2.82 |
| -1.38 | -7.28 | 4.78 | 10 | -3.14 | 0.42 | -0.86 | -7.12 |
| -2.72 | -9.04 | 1.26 | -8.5 | 10 | 4.32 | -1.68 | -1.08 |
| 6.28 | 10 | -3.94 | 0.9 | 4.4 | -0.04 | 1.34 | -1.1 |
| 1.04 | -1.8 | 8.38 | 1.58 | 7 | -6.56 | -1.86 | 1.94 |
| -4.7 | 7.9 | 3.22 | 3.76 | -9.94 | 3.92 | -3.26 | 6.28 |
| -1.88 | -1.06 | 4.04 | -3.28 | -2.38 | -0.48 | 6.2 | 2.26 |
| 2.98 | 3.62 | 2.42 | 4.4 | 4.52 | 6.06 | -5.12 | 4.82 |
| 0.4 | 5.12 | 4.32 | 7.9 | 5.34 | 9.9 | 6.6 | -9.52 |
| 2.6 | 4.28 | -6.3 | 10 | 2.14 | -3.7 | 3.92 | 4.2 |
| -2.64 | 5.933334 | 10 | 10 | 10 | -1.68 | 10 | 4.82 |
| -1.14 | 7.24 | -0.08 | -0.06 | 0.98 | 4.7 | 3.36 | -3.84 |
| 10 | -3.8 | 3.22 | -4.24 | 2.86 | 6.96 | -4.58 | -2.46 |
| -8.18 | 7.82 | -4.38 | -4.62 | -4.84 | 5.78 | 10 | 1.84 |
| -5.84 | -4.54 | 0.72 | 6.02 | 0.14 | 2.14 | 10 | 8.28 |
| 10 | 1.26 | 3.06 | 4.6 | 5.94 | 8.42 | -10 | 5.88 |
| -10 | -5.32 | 5.88 | -5.88 | 5.8 | 10 | -6.32 | 2.84 |

| | | | | | | | |
|---|---|---|---|---|---|---|---|
| -4.86 | -4.06 | -10 | -10 | 3.04 | -6.64 | -9.3 | 7.3 |
| -3.08 | 8.4 | 10 | -8.1 | 1.82 | -8.44 | 5.56 | -10 |
| 4.16 | -5.52 | 10 | -0.26 | -6.56 | -2.38 | 10 | 10 |
| -6.3 | -0.08 | 4.3 | 5.66 | 2.08 | -2.38 | 7.9 | -5.84 |
| 10 | 10 | 10 | 5.52 | -6.32 | -1.06 | 0.84 | -10 |
| -1.9 | -0.54 | -2.64 | 9.64 | 7.14 | -10 | 8.24 | -6.48 |
| -1.54 | -1.82 | 7.6 | -10 | 2.14 | 6.54 | -8.1 | -4.52 |
| 0.54 | 0.38 | 6.54 | 9.78 | 10 | 5.62 | 5.98 | -5.98 |
| 10 | 0.9 | 4.76 | 8.2 | -10 | 4.04 | 1.18 | -7.48 |
| -0.44 | -2.72 | -6.74 | 6 | -5.56 | 7.02 | -7.92 | 0.88 |
| 9.2 | -2.72 | 2.86 | -10 | -7.73333 | 4.08 | 9.4 | 10 |
| 1.86 | -5.36 | 8.34 | 1.78 | 0.66 | 8.56 | 8.26 | -0.42 |
| -4.06 | -3.26 | -5.46 | 3.86 | 10 | -0.32 | 5.46 | 2.78 |
| -0.26 | -0.1 | -2.68 | 2.14 | -8.56 | 8.2 | -6.68 | 4.12 |
| -7.62 | 4.3 | 2.28 | 7.18 | 10 | -4.76 | -5.84 | 2.18 |
| 1.64 | -10 | -5.14 | 2.28 | -4.84 | 7.88 | -4.84 | 0.54 |
| 9.84 | 4.4 | 3.88 | 0.5 | -10 | -8.26 | -1.94 | 1.78 |
| -2.88 | -3.32 | 0.52 | 5.1 | 7.78 | -10 | -10 | -2.92 |
| -10 | 10 | 8.6 | -5.2 | -6.64 | 6.82 | -3.88 | -4.84 |
| 5.82 | 10 | 10 | -3.88 | 2.56 | -5.94 | 4.12 | -4.64 |
| -7.52 | -2.18 | 10 | 6.28 | 2.22 | 8.42 | -3.84 | 2.88 |
| -2.88 | 5.94 | -2.08 | 10 | -0.68 | 4.56 | -4.7 | -1.28 |
| -4.36 | 3.44 | 4.52 | 5 | 10 | 3.46 | 7.8 | 1.04 |
| 7.56 | 0.06 | 9.5 | -10 | 3.82 | 10 | 3.7 | 9.7 |
| -6.26 | 10 | 8.16 | -10 | 4.68 | -3.7 | -7.56 | 1.58 |
| -9.24 | 10 | -8.42 | -5.44 | 10 | -5.06 | 1.24 | -9.84 |
| -8.86 | -5.58 | 10 | -5.44 | 1.62 | -4.04 | -1.96 | -3.2 |
| 0.88 | 10 | -10 | 10 | 4.82 | 6.38 | 3.34 | 4.92 |
| 4.4 | -10 | -4.62 | 10 | 6.72 | 4 | -10 | 0.16 |
| 7.08 | 3.86 | 6.88 | -10 | -1 | 8.5 | 1.34 | 5 |
| 1.14 | 5.84 | 1.21 | 4.48 | 7.1 | -8.82 | -10 | -2.24 |
| -6.44 | -5.26 | 10 | 0.08 | 10 | -3.54 | -2.66 | -2.52 |
| 0.9 | -1.98 | -5.46 | 0.94 | 10 | -2.08 | -2.02 | 0.94 |

| | | | | | | | |
|---|---|---|---|---|---|---|---|
| -4.92 | -4.08 | -3.82 | -5.42 | -10 | -0.76 | -3.26 | -2.9 |
| -1.8 | -3.02 | -6.36 | 5.72 | 7.02 | 7.66 | 10 | 0.48 |
| 0.58 | -0.16 | 6.24 | 5.16 | -9.82 | 5.92 | -2.24 | 6.4 |
| -1.48 | -2 | 3.82 | 1.42 | 2.06 | 4.1 | 7.16 | -4.76 |
| 1.78 | 5.8 | -4.4 | 5.9 | -5.04 | 0.26 | 3.18 | 4.5 |
| -4.08 | -10 | -5.02 | 9.6 | -10 | 5.16 | 1.68 | -2.18 |
| 4.4 | 10 | 4.12 | -9.34 | -8.78 | -1.3 | 1.34 | -10 |
| -4.02 | 5.24 | -4.02 | -2.3 | 1.82 | -3.72 | -10 | -3 |
| 0.38 | -10 | 2.48 | 7.52 | 10 | -5.38 | -8.02 | -0.88 |
| 0.26 | -0.52 | 6.82 | 8.34 | -3.4 | 2.28 | -0.58 | -7.04 |
| 6.1 | -6 | 2.22 | 4.62 | 1.08 | -10 | -10 | -5.8 |
| 10 | 5.86 | -5.96 | 2.52 | 10 | 6.46 | -1.64 | -10 |
| -3.42 | -1.06 | 8.4 | -5.02 | 10 | -10 | -6.12 | 10 |
| 1.76 | 7.86 | 1.5 | 2.44 | 10 | 0.06 | -3.42 | -0.06 |
| 3.58 | -7.68 | -4.6 | 6.54 | -4.98 | 5.38 | 0.5 | -6.24 |
| -3.06 | -1.86 | 2.84 | 3.66 | -7.06 | 10 | -1.36 | -10 |
| 1.96 | -6.18 | -2.06 | 7.7 | 7.4 | 10 | 5.96 | -1.04 |
| 10 | -3.78 | -10 | 10 | -7.41 | -4.46 | 3.64 | -6.8 |
| -3.3 | -4.78 | -4.82 | 7.1 | 6.26 | 0.62 | -0.9 | -10 |
| 8.48 | -4.08 | -1.46 | -1.98 | -3.48 | 0.94 | 6.76 | -10 |
| -4.42 | 3.88 | 7.24 | 4.32 | 10 | -4.22 | -1.92 | 4.14 |
| 1.56 | 10 | -4.66 | 2.9 | 8.62 | -3 | -4.42 | -6.42 |
| -8.2 | -6.42 | -0.32 | 3.5 | 10 | 7.3 | -4.3 | 6.46 |
| 10 | -2.76 | 8.1 | 0.24 | 10 | 10 | 7.58 | 4.24 |
| 8.16 | -3.46 | -6.28 | -2.42 | 3.82 | 3.86 | 9.96 | 3.32 |
| 10 | -8.36 | -3.52 | -7.72 | -5.84 | 6.76 | 8.76 | -1.72 |
| -0.28 | -5.24 | -4.68 | 8.76 | 1.924 | -0.1 | -10 | -8.96 |
| 5.12 | 10 | 2.8 | 6.7 | 3.52 | -8.22 | 2.76 | 0.64 |
| 5.22 | 10 | 8.08 | 3.9 | -5.62 | 5.38 | -2.44 | 6.58 |
| -3.62 | 4.24 | 1.94 | 10 | -1.22 | 7.96 | 7.62 | -1.88 |
| 0.56 | -8.14 | -2.16 | 0.24 | -5.54 | 8.1 | -1.96 | 9.28 |
| -1.9 | -2.72 | 8.9 | 8.46 | 9.5 | 6.76 | -3.5 | 10 |
| -0.54 | 9.12 | 4.62 | -2.98 | 6.66 | 6.08 | 2.58 | -7.62 |

| | | | | | | | |
|---|---|---|---|---|---|---|---|
| -6.46 | 2.06 | 10 | -7.02 | -0.16 | -10 | 6.3 | 5.16 |
| 0.32 | 1.4 | -4.26 | 10 | 3.38 | -5.8 | -0.2 | -10 |
| 3.16 | -2.14 | -3.22 | 9.18 | 0.32 | 6.3 | 2.14 | 7.48 |
| | 4.1 | 5.22 | 4.68 | 3.2 | 10 | 10 | 4.62 |
| | -1.92 | 4.58 | 10 | 10 | 7.62 | 10 | -10 |
| | -3.92 | -6.04 | -2.16 | 2.02 | 3.04 | 5.16 | 8.26 |
| | 7.66 | 8.24 | 5.78 | -8.2 | 10 | 0.46 | 2.02 |
| | -1.8 | -6.04 | 10 | 2.94 | 10 | 10 | |
| | -5.82 | 0.16 | 10 | -2.06 | 6 | -0.06 | |
| | 0.84 | 6.48 | -8.12 | -0.98 | 6.04 | 10 | |
| | 7.78 | -4.52 | 2.44 | | 3.22 | 6.32 | |
| | | -0.06 | -2.94 | | 3.12 | | |
| | | -0.08 | -10 | | -5.88 | | |
| | | -5.98 | -9.84 | | 0.62 | | |
| | | 7.92 | -0.82 | | 8.32 | | |
| | | 2.28 | -1.72 | | 6.74 | | |
| | | 5.98 | -3.76 | | -10 | | |
| | | -3.12 | -0.14 | | -7.7 | | |
| | | 4.1 | -3.1 | | -3.46 | | |
| | | -2.16 | 9.14 | | 3.24 | | |
| | | 1.98 | -1.32 | | 0.66 | | |
| | | -3.58 | -1.86 | | | | |
| | | 2.06 | -0.06 | | | | |
| | | -6.48 | -5.94 | | | | |
| | | 0.42 | -3.34 | | | | |
| | | 4.9 | 2.36 | | | | |
| | | 10 | 1.72 | | | | |
| | | 6.02 | 7.5 | | | | |
| | | -4.6 | 10 | | | | |
| | | 9.92 | 6.04 | | | | |
| | | -9.58 | | | | | |
| | | 9.02 | | | | | |
| | | 6.92 | | | | | |

**RAW DATA**

| 0.1 ng/ul | CONTROL | 0.5 ng/ul | CONTROL | 0.9 ng/ul | CONTROL | 1.3 ng/ul | CONTROL |
|---|---|---|---|---|---|---|---|
| 1.39 | 8.61 | 3.97 | 6.03 | 5.38 | 4.62 | 1.97 | 8.03 |
| 1.07 | 8.93 | 4.82 | 5.18 | 6.71 | 3.29 | 6.23 | 3.77 |
| 4.27 | 5.73 | 6.86 | 3.14 | 5.84 | 4.16 | 6.27 | 3.73 |
| 3.91 | 6.09 | 6.95 | 3.05 | 6.4 | 3.6 | 5.93 | 4.07 |
| 2.05 | 7.95 | 6.38 | 3.62 | 2.3 | 7.7 | 2.25 | 7.75 |
| 5.05 | 4.95 | 5.71 | 4.29 | 0 | 10 | 1.75 | 8.25 |
| 1.87 | 8.13 | 5.11 | 4.89 | 7.02 | 2.98 | 1.39 | 8.61 |
| 4.56 | 5.44 | 7.55 | 2.45 | 0 | 10 | 0.37 | 9.63 |
| 5.83 | 4.17 | 3.55 | 6.45 | 5.05 | 4.95 | 2.22 | 7.78 |
| 0.68 | 9.32 | 3.24 | 6.76 | 3.86 | 6.14 | 9.5 | 0.5 |
| 5.42 | 4.58 | 6.66 | 3.34 | 3.82 | 6.18 | 10 | 0 |
| 4.37 | 5.63 | 6.02 | 3.98 | 6.42 | 3.58 | 8.7 | 1.3 |
| 7.65 | 2.35 | 10 | 0 | 4.52 | 5.48 | 8.76 | 1.24 |
| 5.18 | 4.82 | 1.23 | 8.77 | 1.21 | 8.79 | 1.8 | 8.2 |
| 5.69 | 4.31 | 8.64 | 1.36 | 2.61 | 7.39 | 0 | 10 |
| 6.36 | 3.64 | 9.52 | 0.48 | 4.37 | 5.63 | 9.25 | 0.75 |
| 1.86 | 8.14 | 0 | 10 | 6.97 | 3.03 | 4.55 | 5.45 |
| 4.48 | 5.52 | 5.9 | 4.1 | 0.81 | 9.19 | 4.21 | 5.79 |
| 7.35 | 2.65 | 1.05 | 8.95 | 3.39 | 6.61 | 3.12 | 6.88 |
| 5.94 | 4.06 | 5.53 | 4.47 | 2.98 | 7.02 | 6.64 | 3.36 |
| 3.51 | 6.49 | 3.19 | 6.81 | 3.79 | 6.21 | 2.8 | 7.2 |
| 4.8 | 5.2 | 2.44 | 7.56 | 2.84 | 7.16 | 1.05 | 8.95 |
| 3.7 | 6.3 | 2.86 | 7.14 | 8.15 | 1.85 | 0 | 10 |
| 6.32 | 3.68 | 2.033333 | 7.966667 | 0 | 10 | 0 | 10 |
| 5.57 | 4.43 | 1.38 | 8.62 | 5.04 | 4.96 | 5.03 | 4.97 |
| 0 | 10 | 6.9 | 3.1 | 3.39 | 6.61 | 7.12 | 2.88 |
| 9.09 | 0.91 | 1.09 | 8.91 | 7.19 | 2.81 | 7.31 | 2.69 |
| 7.92 | 2.08 | 7.27 | 2.73 | 4.64 | 5.36 | 1.99 | 8.01 |

| | | | | | | | |
|---|---|---|---|---|---|---|---|
| 0 | 10 | 4.37 | 5.63 | 3.47 | 6.53 | 2.7 | 7.3 |
| 10 | 0 | 7.66 | 2.34 | 2.06 | 7.94 | 7.94 | 2.06 |
| 7.43 | 2.57 | 7.03 | 2.97 | 10 | 0 | 10 | 0 |
| 6.54 | 3.46 | 0.8 | 9.2 | 0 | 10 | 9.05 | 0.95 |
| 2.92 | 7.08 | 7.76 | 2.24 | 0 | 10 | 5.13 | 4.87 |
| 8.15 | 1.85 | 5.04 | 4.96 | 2.85 | 7.15 | 2.17 | 7.83 |
| 0 | 10 | 0 | 10 | 0 | 10 | 2.24 | 7.76 |
| 5.95 | 4.05 | 5.27 | 4.73 | 6.32 | 3.68 | 0.18 | 9.82 |
| 5.77 | 4.23 | 5.91 | 4.09 | 1.2 | 8.8 | 10 | 0 |
| 4.73 | 5.27 | 4.81 | 5.19 | 1.73 | 8.27 | 0.11 | 9.89 |
| 0 | 10 | 4.55 | 5.45 | 2.62 | 7.38 | 0.9 | 9.1 |
| 5.22 | 4.78 | 6.36 | 3.64 | 8.37 | 1.63 | 2 | 8 |
| 0.4 | 9.6 | 6.36 | 3.64 | 3.57 | 6.43 | 10 | 0 |
| 4.07 | 5.93 | 7.68 | 2.32 | 0.83 | 9.17 | 4.11 | 5.89 |
| 7.03 | 2.97 | 6.63 | 3.37 | 7.73 | 2.27 | 3.07 | 6.93 |
| 5.13 | 4.87 | 5.05 | 4.95 | 6.34 | 3.66 | 3.93 | 6.07 |
| 8.81 | 1.19 | 2.85 | 7.15 | 3.86 | 6.14 | 1.41 | 8.59 |
| 4.18 | 5.82 | 10 | 0 | 7.57 | 2.43 | 3.86 | 6.14 |
| 0.08 | 9.92 | 2.8 | 7.2 | 3.06 | 6.94 | 4.75 | 5.25 |
| 6.44 | 3.56 | 6.66 | 3.34 | 4.74 | 5.26 | 2.45 | 7.55 |
| 10 | 0 | 0 | 10 | 0.7 | 9.3 | 7.6 | 2.4 |
| 2.09 | 7.91 | 0 | 10 | 0 | 10 | 6.94 | 3.06 |
| 8.76 | 1.24 | 6.09 | 3.91 | 0 | 10 | 1.86 | 8.14 |
| 6.44 | 3.56 | 2.03 | 7.97 | 6.04 | 3.96 | 0 | 10 |
| 7.18 | 2.82 | 3.28 | 6.72 | 2.74 | 7.26 | 2.5 | 7.5 |
| 1.22 | 8.78 | 4.97 | 5.03 | 0.25 | 9.75 | 10 | 0 |
| 8.13 | 1.87 | 0 | 10 | 0.92 | 9.08 | 10 | 0 |
| 9.62 | 0.38 | 0 | 10 | 9.21 | 0.79 | 7.72 | 2.28 |
| 9.43 | 0.57 | 7.79 | 2.21 | 0 | 10 | 7.72 | 2.28 |
| 4.56 | 5.44 | 0 | 10 | 10 | 0 | 0 | 10 |
| 2.8 | 7.2 | 10 | 0 | 7.31 | 2.69 | 0 | 10 |
| 1.46 | 8.54 | 3.07 | 6.93 | 1.56 | 8.44 | 10 | 0 |
| 4.43 | 5.57 | 2.08 | 7.92 | 4.38 | 5.59 | 2.76 | 7.24 |

| | | | | | | | |
|---|---|---|---|---|---|---|---|
| 8.22 | 1.78 | 7.63 | 2.37 | 0 | 10 | 4.96 | 5.04 |
| 4.55 | 5.45 | 5.99 | 4.01 | 7.73 | 2.27 | 4.53 | 5.47 |
| 7.46 | 2.54 | 7.04 | 2.96 | 6.91 | 3.09 | 7.71 | 2.29 |
| 5.9 | 4.1 | 6.51 | 3.49 | 8.18 | 1.82 | 2.14 | 7.86 |
| 4.71 | 5.29 | 5.08 | 4.92 | 1.88 | 8.12 | 2.42 | 7.58 |
| 5.74 | 4.26 | 6 | 4 | 3.09 | 6.91 | 4.29 | 5.71 |
| 4.11 | 5.89 | 2.1 | 7.9 | 7.2 | 2.8 | 2.05 | 7.95 |
| 7.04 | 2.96 | 10 | 0 | 7.51 | 2.49 | 0.2 | 9.8 |
| 2.8 | 7.2 | 0 | 10 | 2.94 | 7.06 | 9.67 | 0.33 |
| 7.01 | 2.99 | 2.38 | 7.62 | 7.01 | 2.99 | 6.15 | 3.85 |
| 4.81 | 5.19 | 10 | 0 | 3.76 | 6.24 | 1.24 | 8.76 |
| 4.87 | 5.13 | 5.26 | 4.74 | 1.59 | 8.41 | 0.83 | 9.17 |
| 1.95 | 8.05 | 8 | 2 | 3.89 | 6.11 | 2.69 | 7.31 |
| 0 | 10 | 2.07 | 7.93 | 7.98 | 2.02 | 3.74 | 6.26 |
| 6.71 | 3.29 | 5.53 | 4.47 | 0.8 | 9.2 | 7.51 | 2.49 |
| 4.12 | 5.88 | 1.07 | 8.93 | 4.25 | 5.75 | 3.78 | 6.22 |
| 3.21 | 6.79 | 8.84 | 1.16 | 7.3 | 2.7 | 1.73 | 8.27 |
| 6.53 | 3.47 | 5.93 | 4.07 | 3.58 | 6.42 | 3.17 | 6.83 |
| 4.02 | 5.98 | 8.09 | 1.91 | 6.03 | 3.97 | 1.15 | 8.85 |
| 0 | 10 | 6.89 | 3.11 | 10 | 0 | 0 | 10 |
| 6.65 | 3.35 | 7.39 | 2.61 | 7.41 | 2.59 | 1.45 | 8.55 |
| 0.76 | 9.24 | 7.04 | 2.96 | 5.73 | 4.27 | 5.99 | 4.01 |
| 7.21 | 2.79 | 3.06 | 6.94 | 1.38 | 8.62 | 2.84 | 7.16 |
| 4.22 | 5.78 | 0 | 10 | 7.33 | 2.67 | 3.55 | 6.45 |
| 9.1 | 0.9 | 8.21 | 1.79 | 5.16 | 4.84 | 3.25 | 6.75 |
| 0 | 10 | 6.38 | 3.62 | 0.95 | 9.05 | 4.88 | 5.12 |
| 0.92 | 9.08 | 6.73 | 3.27 | 8.14 | 1.86 | 6.21 | 3.79 |
| 0 | 10 | 9.18 | 0.82 | 6.76 | 3.24 | 8.86 | 1.14 |
| 5.14 | 4.86 | 7.62 | 2.38 | 7.34 | 2.66 | 0.62 | 9.38 |
| 2.44 | 7.56 | 0 | 10 | 3.6 | 6.4 | 1.65 | 8.35 |
| 2.39 | 7.61 | 0 | 10 | 0.96 | 9.04 | 3.05 | 6.95 |
| 6.81 | 3.19 | 2.88 | 7.12 | 4.03 | 5.97 | 0 | 10 |
| 4.72 | 5.28 | 9.07 | 0.93 | 6.08 | 3.92 | 4.88 | 5.12 |

| | | | | | | | |
|---|---|---|---|---|---|---|---|
| 5.95 | 4.05 | 6.36 | 3.64 | 0.55 | 9.45 | 0.77 | 9.23 |
| 5.27 | 4.73 | 0.44 | 9.56 | 2.69 | 7.31 | 6.49 | 3.51 |
| 8.23 | 1.77 | 3.97 | 6.03 | 0 | 10 | 8.51 | 1.49 |
| 4.84 | 5.16 | 4.3 | 5.7 | 7.13 | 2.87 | 0 | 10 |
| 3.42 | 6.58 | 6.07 | 3.93 | 6.61 | 3.39 | 0.41 | 9.59 |
| | | 2.95 | 7.05 | 2.39 | 7.61 | 2.66 | 7.34 |
| | | 5.96 | 4.04 | 2.71 | 7.29 | 0 | 10 |
| | | 6.96 | 3.04 | 8.02 | 1.98 | 6.08 | 3.92 |
| | | 1.17 | 8.83 | 0.88 | 9.12 | 2.11 | 7.89 |
| | | 5.9 | 4.1 | 8.02 | 1.98 | 0 | 10 |
| | | 7.91 | 2.09 | 4.92 | 5.08 | 0 | 10 |
| | | 4.58 | 5.42 | 1.76 | 8.24 | 9.06 | 0.94 |
| | | 1.11 | 8.89 | 7.26 | 2.74 | 3.78 | 6.22 |
| | | | | 5.03 | 4.97 | 6.47 | 3.53 |
| | | | | 5.04 | 4.96 | 10 | 0 |
| | | | | 7.99 | 2.01 | 9.92 | 0.08 |
| | | | | 1.04 | 8.96 | 5.41 | 4.59 |
| | | | | 3.86 | 6.14 | 5.86 | 4.14 |
| | | | | 2.01 | 7.99 | 6.88 | 3.12 |
| | | | | 6.56 | 3.44 | 5.07 | 4.93 |
| | | | | 2.95 | 7.05 | 6.55 | 3.45 |
| | | | | 6.08 | 3.92 | 0.43 | 9.57 |
| | | | | 4.01 | 5.99 | 5.66 | 4.34 |
| | | | | 6.79 | 3.21 | 5.93 | 4.07 |
| | | | | 3.97 | 6.03 | 5.03 | 4.97 |
| | | | | 8.24 | 1.76 | 7.97 | 2.03 |
| | | | | 4.79 | 5.21 | 6.67 | 3.33 |
| | | | | 2.55 | 7.45 | 3.82 | 6.18 |
| | | | | 0 | 10 | 4.14 | 5.86 |
| | | | | 1.99 | 8.01 | 1.25 | 8.75 |
| | | | | 7.3 | 2.7 | 0 | 10 |
| | | | | 0.04 | 9.96 | 1.98 | 8.02 |
| | | | | 9.79 | 0.21 | | |

| | | | | | |
|---|---|---|---|---|---|
| | | | | 0.49 | 9.51 |
| | | | | 1.54 | 8.46 |

| 1.7 ng/ul | CONTROL | 2.1 ng/ul | CONTROL | 2.5 ng/ul | CONTROL | 2.9 ng/ul | CONTROL |
|---|---|---|---|---|---|---|---|
| 2.49 | 7.51 | 7.2 | 2.8 | 8.1 | 1.9 | 1.92 | 8.08 |
| 6.55 | 3.45 | 6.87 | 3.13 | 3.1 | 6.9 | 0 | 10 |
| 6.79 | 3.21 | 8.44 | 1.56 | 0 | 10 | 3.5 | 6.5 |
| 10 | 0 | 4.39 | 5.61 | 3.71 | 6.29 | 7.63 | 2.37 |
| 10 | 0 | 10 | 0 | 9.2 | 0.8 | 7.91 | 2.09 |
| 4.25 | 5.75 | 4.99 | 5.01 | 8.91 | 1.09 | 0 | 10 |
| 2.29 | 7.71 | 1.17 | 8.83 | 3.99 | 6.01 | 3.06 | 6.94 |
| 3.12 | 6.88 | 8.06 | 1.94 | 5.5 | 4.5 | 9.23 | 0.77 |
| 0.68 | 9.32 | 10 | 0 | 1.02 | 8.98 | 8.55 | 1.45 |
| 6.71 | 3.29 | 10 | 0 | 2.17 | 7.83 | 7.68 | 2.32 |
| 0 | 10 | 2.1 | 7.9 | 8.01 | 1.99 | 2.67 | 7.33 |
| 0 | 10 | 4.92 | 5.08 | 4.82 | 5.18 | 0.06 | 9.94 |
| 7.03 | 2.97 | 1.6 | 8.4 | 8.37 | 1.63 | 7.97 | 2.03 |
| 5.83 | 4.17 | 1.93 | 8.07 | 0.93 | 9.07 | 3.59 | 6.41 |
| 6.57 | 3.43 | 4.79 | 5.21 | 5.43 | 4.57 | 8.56 | 1.44 |
| 0 | 10 | 2.84 | 7.16 | 5.84 | 4.16 | 5.54 | 4.46 |
| 2.8 | 7.2 | 5.02 | 4.98 | 4.33 | 5.67 | 5.55 | 4.45 |
| 1.5 | 8.5 | 8.28 | 1.72 | 5.93 | 4.07 | 4.03 | 5.97 |
| 9.97 | 0.03 | 3.04 | 6.96 | 6.63 | 3.37 | 1.86 | 8.14 |
| 6.19 | 3.81 | 5.24 | 4.76 | 1.9 | 8.1 | 3.87 | 6.13 |
| 2.74 | 7.26 | 1.97 | 8.03 | 7.56 | 2.44 | 2.59 | 7.41 |
| 2.33 | 7.67 | 0.05 | 9.95 | 1.7 | 8.3 | 9.76 | 0.24 |
| 3.93 | 6.07 | 6.85 | 3.15 | 3.04 | 6.96 | 2.9 | 7.1 |
| 0 | 10 | 5.84 | 4.16 | 0 | 10 | 2.59 | 7.41 |
| 4.51 | 5.49 | 2.65 | 7.35 | 3.32 | 6.68 | 6.92 | 3.08 |
| 3.57 | 6.43 | 1.52 | 8.48 | 7.29 | 2.71 | 6.23 | 3.77 |
| 7.42 | 2.58 | 2.11 | 7.89 | 0 | 10 | 4.08 | 5.92 |
| 4.93 | 5.07 | 3.93 | 6.07 | 0 | 10 | 0.86 | 9.14 |
| 2.03 | 7.97 | 0.79 | 9.21 | 10 | 0 | 2.06 | 7.94 |

| | | | | | | | |
|---|---|---|---|---|---|---|---|
| 2.1 | 7.9 | 0 | 10 | 8.16 | 1.84 | 3.58 | 6.42 |
| 3.48 | 6.52 | 8.32 | 1.68 | 9.65 | 0.35 | 1.35 | 8.65 |
| 4.09 | 5.91 | 9.22 | 0.78 | 2.22 | 7.78 | 10 | 0 |
| 8.28 | 1.72 | 6.19 | 3.81 | 0 | 10 | 0 | 10 |
| 3.96 | 6.04 | 6.19 | 3.81 | 1.05 | 8.95 | 7.92 | 2.08 |
| 8.16 | 1.84 | 5.53 | 4.47 | 4.58 | 5.42 | 10 | 0 |
| 1.93 | 9.07 | 10 | 0 | 0.88 | 9.12 | 8.24 | 1.76 |
| 3.93 | 6.07 | 1.73 | 8.27 | 9.05 | 0.95 | 7.26 | 2.74 |
| 0 | 10 | 2.19 | 7.81 | 2.01 | 7.99 | 7.99 | 2.01 |
| 10 | 0 | 2.98 | 7.02 | 4.41 | 5.59 | 8.74 | 1.26 |
| 7.78 | 2.22 | 1.49 | 8.51 | 8.96 | 1.04 | 4.56 | 5.44 |
| 8.866667 | 1.133333 | 2.96 | 7.04 | 0.3 | 9.7 | 0 | 10 |
| 4.67 | 5.33 | 0.72 | 9.28 | 0.87 | 9.13 | 5.21 | 4.79 |
| 0 | 10 | 5.16 | 4.84 | 2.27 | 7.73 | 3.61 | 6.39 |
| 9.28 | 0.72 | 0.9 | 9.1 | 8.34 | 1.66 | 2.94 | 7.06 |
| 0 | 10 | 7.38 | 2.62 | 7.92 | 2.08 | 3.91 | 6.09 |
| 7.42 | 2.58 | 1.06 | 8.94 | 7.42 | 2.58 | 4.73 | 5.27 |
| 10 | 0 | 9.13 | 0.87 | 5.97 | 4.03 | 4.11 | 5.89 |
| 1.11 | 8.89 | 10 | 0 | 10 | 0 | 6.46 | 3.54 |
| 8.32 | 1.68 | 1.59 | 8.41 | 6.94 | 3.06 | 7.42 | 2.58 |
| 3.72 | 6.28 | 7.97 | 2.03 | 2.94 | 7.06 | 7.32 | 2.68 |
| 3.89 | 6.11 | 0.79 | 9.21 | 6.92 | 3.08 | 3.56 | 6.44 |
| 5.34 | 4.66 | 2.72 | 7.28 | 7.35 | 2.65 | 5.64 | 4.36 |
| 0 | 10 | 3.27 | 6.73 | 1.1 | 8.9 | 4.48 | 5.52 |
| 3.09 | 6.91 | 0 | 10 | 3.15 | 6.85 | 0.15 | 9.85 |
| 2.66 | 7.34 | 6.85 | 3.15 | 8.78 | 1.22 | 4.21 | 5.79 |
| 0 | 10 | 7.53 | 2.47 | 4.38 | 5.62 | 9.92 | 0.08 |
| 4.19 | 5.81 | 7.02 | 2.98 | 5.98 | 4.02 | 6.6 | 3.4 |
| 2.59 | 7.41 | 1.81 | 8.19 | 3.33 | 6.67 | 2.54 | 7.46 |
| 1.64 | 8.36 | 3 | 7 | 10 | 0 | 4.92 | 5.08 |
| 5.5 | 4.5 | 0.75 | 9.25 | 4.33 | 5.67 | 2.5 | 7.5 |
| 1.45 | 8.55 | 9.41 | 0.59 | 10 | 0 | 6.12 | 3.88 |
| 0 | 10 | 6.77 | 3.23 | 6.33 | 3.67 | 6.26 | 3.74 |

| | | | | | | | |
|---|---|---|---|---|---|---|---|
| 0 | 10 | 6.04 | 3.96 | 6.01 | 3.99 | 4.53 | 5.47 |
| 10 | 0 | 5.38 | 4.62 | 6.63 | 3.37 | 6.45 | 3.55 |
| 1.49 | 8.51 | 1.17 | 8.83 | 0 | 10 | 4.76 | 5.24 |
| 9.91 | 0.09 | 2.04 | 7.96 | 6.12 | 3.88 | 1.8 | 8.2 |
| 3.97 | 6.03 | 2.95 | 7.05 | 1.42 | 8.58 | 7.38 | 2.62 |
| 7.52 | 2.48 | 4.87 | 5.13 | 3.41 | 6.59 | 2.75 | 7.25 |
| 10 | 0 | 2.42 | 7.58 | 4.16 | 5.84 | 6.09 | 3.91 |
| 9.39 | 0.61 | 5.65 | 4.35 | 4.33 | 5.67 | 10 | 0 |
| 4.09 | 5.91 | 6.86 | 3.14 | 10 | 0 | 6.5 | 3.5 |
| 0 | 10 | 7.69 | 2.31 | 9.01 | 0.99 | 5.44 | 4.56 |
| 6.7 | 3.3 | 3.86 | 6.14 | 5.29 | 4.71 | 8.52 | 1.48 |
| 4.46 | 5.54 | 10 | 0 | 10 | 0 | 7.9 | 2.1 |
| 0 | 10 | 1.77 | 8.23 | 5.82 | 4.18 | 10 | 0 |
| 0 | 10 | 10 | 0 | 8.06 | 1.94 | 0 | 10 |
| 0 | 10 | 4.97 | 5.03 | 6.71 | 3.29 | 5.03 | 4.97 |
| 7.49 | 2.51 | 2.31 | 7.69 | 4.75 | 5.25 | 8.12 | 1.88 |
| 8.53 | 1.47 | 0 | 10 | 5.68 | 4.32 | 10 | 0 |
| 1.3 | 8.7 | 0 | 10 | 2.02 | 7.98 | 5.52 | 4.48 |
| 8.44 | 1.03 | 7.23 | 2.77 | 3.18 | 6.82 | 8.4 | 1.6 |
| 1.87 | 8.13 | 4.69 | 5.31 | 5.45 | 4.55 | 10 | 0 |
| 6.74 | 3.26 | 4.53 | 5.47 | 1.62 | 8.38 | 10 | 0 |
| 0 | 10 | 7.11 | 2.89 | 5.96 | 4.04 | 2.93 | 7.07 |
| 0.69 | 9.31 | 6.5 | 3.5 | 7.21 | 2.79 | 8.21 | 1.79 |
| 0 | 10 | 1.35 | 8.65 | 7.15 | 2.85 | 1.77 | 8.23 |
| 0 | 10 | 0 | 10 | 1.21 | 8.79 | 2.88 | 7.12 |
| 3.09 | 6.91 | 3.07 | 6.93 | 0.02 | 9.98 | 3.34 | 6.66 |
| 7.92 | 2.08 | 1.62 | 8.38 | 0.62 | 9.38 | 5.86 | 4.14 |
| 4.038 | 5.962 | 5.05 | 4.95 | 10 | 0 | 9.48 | 0.52 |
| 3.24 | 6.76 | 9.11 | 0.89 | 3.62 | 6.38 | 4.68 | 5.32 |
| 7.81 | 2.19 | 2.31 | 7.69 | 6.22 | 3.78 | 1.71 | 8.29 |
| 5.61 | 4.39 | 1.02 | 8.98 | 1.19 | 8.81 | 5.94 | 4.06 |
| 7.77 | 2.23 | 0.95 | 9.05 | 5.98 | 4.02 | 0.36 | 9.64 |
| 0.25 | 9.75 | 1.62 | 8.38 | 6.75 | 3.25 | 0 | 10 |

| | | | | | | | |
|---|---|---|---|---|---|---|---|
| 1.67 | 8.33 | 1.96 | 8.04 | 3.71 | 6.29 | 8.81 | 1.19 |
| 5.08 | 4.92 | 10 | 0 | 1.85 | 8.15 | 2.42 | 7.58 |
| 3.31 | 6.69 | 7.9 | 2.1 | 5.1 | 4.9 | 10 | 0 |
| 4.84 | 5.16 | 1.85 | 8.15 | 3.93 | 6.07 | 1.26 | 8.74 |
| 3.4 | 6.6 | 0 | 10 | 0 | 10 | 2.69 | 7.31 |
| 0 | 10 | 1.19 | 8.81 | 0 | 10 | 10 | 0 |
| 3.99 | 6.01 | 3.48 | 6.52 | 2.42 | 7.58 | 0.87 | 9.13 |
| 9.1 | 0.9 | 0 | 10 | 4.77 | 5.23 | 3.99 | 6.01 |
| 3.53 | 6.47 | 0 | 10 | 0 | 10 | | |
| 6.03 | 3.97 | 2 | 8 | 5.03 | 4.97 | | |
| 5.49 | 4.51 | 1.98 | 8.02 | 0 | 10 | | |
| | | 3.39 | 6.61 | 1.84 | 8.16 | | |
| | | 3.44 | 6.56 | | | | |
| | | 7.94 | 2.06 | | | | |
| | | 4.69 | 5.31 | | | | |
| | | 0.84 | 9.16 | | | | |
| | | 1.63 | 8.37 | | | | |
| | | 10 | 0 | | | | |
| | | 8.85 | 1.15 | | | | |
| | | 6.73 | 3.27 | | | | |
| | | 3.38 | 6.62 | | | | |
| | | 4.67 | 5.33 | | | | |

**Figure 5**

**Caryophyllene 0.17 ug/ul +Copaene 1.7 ng/ul**

| Treatment | Control |
|---|---|
| 2.53 | 7.47 |
| 8.63 | 1.37 |
| 2.45 | 7.55 |
| 10 | 0 |
| 7.8 | 2.2 |
| 5.95 | 4.05 |
| 2.84 | 7.16 |
| 6.56 | 3.44 |
| 10 | 0 |
| 1.07 | 8.93 |
| 4.83 | 5.17 |
| 7.3 | 2.7 |
| 8.1 | 1.9 |
| 4.03 | 5.97 |
| 0 | 10 |
| 0 | 10 |
| 3.95 | 6.05 |
| 3.71 | 6.29 |
| 0 | 10 |
| 2.82 | 7.18 |
| 4.14 | 5.86 |
| 1.95 | 8.05 |
| 4 | 6 |
| 10 | 0 |
| 4.02 | 5.98 |
| 9.95 | 0.05 |
| 1.22 | 8.78 |
| 9.68 | 0.32 |
| 10 | 0 |
| 2.85 | 7.15 |
| 0 | 10 |
| 0 | 10 |
| 5.22 | 4.78 |
| 2.84 | 7.16 |
| 4.11 | 5.89 |
| 10 | 0 |
| 10 | 0 |
| 7.55 | 2.45 |
| 1.5 | 8.5 |
| 8.77 | 1.23 |
| 5.07 | 4.93 |

**Caryophyllene 0.13 ug/ul +Copaene 1.3 ng/ul**

| Treatment | Control |
|---|---|
| 9.75 | 0.25 |
| 9.39 | 0.61 |
| 9.23 | 0.77 |
| 9 | 1 |
| 8.98 | 1.02 |
| 8.78 | 1.22 |
| 8.49 | 1.51 |
| 8.45 | 1.55 |
| 8.28 | 1.72 |
| 8.02 | 1.98 |
| 7.91 | 2.09 |
| 7.89 | 2.11 |
| 7.84 | 2.16 |
| 7.73 | 2.27 |
| 7.7 | 2.3 |
| 7.5 | 2.5 |
| 7.46 | 2.54 |
| 7.31 | 2.69 |
| 7.2 | 2.8 |
| 7.02 | 2.98 |
| 7.01 | 2.99 |
| 6.71 | 3.29 |
| 6.7 | 3.3 |
| 6.61 | 3.39 |
| 6.6 | 3.4 |
| 6.34 | 3.66 |
| 6.29 | 3.71 |
| 6.2 | 3.8 |
| 6.18 | 3.82 |
| 6.1 | 3.9 |
| 6.05 | 3.95 |
| 6.02 | 3.98 |
| 6 | 4 |
| 5.84 | 4.16 |
| 5.73 | 4.27 |
| 5.68 | 4.32 |
| 5.57 | 4.43 |
| 5.54 | 4.46 |
| 5.53 | 4.47 |
| 5.44 | 4.56 |
| 5.22 | 4.78 |

| | |
|---|---|
| 4.86 | 5.14 |
| 2.12 | 7.88 |
| 2.96 | 7.04 |
| 9.62 | 0.38 |
| 0 | 10 |
| 2.7 | 7.3 |
| 4.6 | 5.4 |
| 4.98 | 5.02 |
| 9.13 | 0.87 |
| 4.55 | 5.45 |
| 4.37 | 5.63 |
| 3.88 | 6.12 |
| 0.38 | 9.62 |
| 0 | 10 |
| 5.08 | 4.92 |
| 8.07 | 1.93 |
| 3.11 | 6.89 |
| 6.03 | 3.97 |
| 5.24 | 4.76 |
| 4.11 | 5.89 |
| 5.38 | 4.62 |
| 7.59 | 2.41 |
| 10 | 0 |
| 3.58 | 6.42 |
| 3.57 | 6.43 |
| 4.1 | 5.9 |
| 6.9 | 3.1 |
| 5.77 | 4.23 |
| 4.51 | 5.49 |
| 8.86 | 1.14 |
| 8.59 | 1.41 |
| 8.34 | 1.66 |
| 4.23 | 5.77 |
| 6.47 | 3.53 |
| 6.19 | 3.81 |
| 4.59 | 5.41 |
| 6.08 | 3.92 |
| 9.36 | 0.64 |
| 1.96 | 8.04 |
| 10 | 0 |
| 6.99 | 3.01 |
| 2.82 | 7.18 |
| 3.8 | 6.2 |
| 7.48 | 2.52 |
| 6.68 | 3.32 |

| | |
|---|---|
| 5.12 | 4.88 |
| 5.05 | 4.95 |
| 4.92 | 5.08 |
| 4.83 | 5.17 |
| 4.77 | 5.23 |
| 4.73 | 5.27 |
| 4.73 | 5.27 |
| 4.63 | 5.37 |
| 4.58 | 5.42 |
| 4.52 | 5.48 |
| 4.46 | 5.54 |
| 4.34 | 5.66 |
| 4.29 | 5.71 |
| 4.21 | 5.79 |
| 4.13 | 5.87 |
| 4.04 | 5.96 |
| 4.01 | 5.99 |
| 3.96 | 6.04 |
| 3.89 | 6.11 |
| 3.86 | 6.14 |
| 3.81 | 6.19 |
| 3.8 | 6.2 |
| 3.73 | 6.27 |
| 3.71 | 6.29 |
| 3.68 | 6.32 |
| 3.65 | 6.35 |
| 3.63 | 6.37 |
| 3.61 | 6.39 |
| 3.6 | 6.4 |
| 3.56 | 6.44 |
| 3.52 | 6.48 |
| 3.29 | 6.71 |
| 3.14 | 6.86 |
| 3.05 | 6.95 |
| 3.02 | 6.98 |
| 2.98 | 7.02 |
| 2.98 | 7.02 |
| 2.97 | 7.03 |
| 2.91 | 7.09 |
| 2.84 | 7.16 |
| 2.75 | 7.25 |
| 2.64 | 7.36 |
| 2.59 | 7.41 |
| 2.44 | 7.56 |
| 2.25 | 7.75 |

| | |
|---|---|
| 5.67 | 4.33 |
| 0 | 10 |
| 3.76 | 6.24 |
| 2.83 | 7.17 |
| 7.41 | 2.59 |
| 8.1 | 1.9 |
| 5.99 | 4.01 |
| 5.62 | 4.38 |
| 8.01 | 1.99 |
| 3.93 | 6.07 |
| 0 | 10 |
| 4.42 | 5.58 |
| 7.59 | 2.41 |
| 6.69 | 3.31 |

| | |
|---|---|
| 2.24 | 7.76 |
| 2.16 | 7.84 |
| 2.12 | 7.88 |
| 2.07 | 7.93 |
| 2.03 | 7.97 |
| 1.93 | 8.07 |
| 1.55 | 8.45 |
| 1.46 | 8.54 |
| 1.42 | 8.58 |
| 1.36 | 8.64 |
| 1.25 | 8.75 |
| 1.22 | 8.78 |
| 1.21 | 8.79 |
| 1.12 | 8.88 |
| 1.02 | 8.98 |
| 0.97 | 9.03 |
| 0.85 | 9.15 |
| 0.53 | 9.47 |
| 0.52 | 9.48 |
| 0.48 | 9.52 |
| 0.12 | 9.88 |
| 0.07 | 9.93 |
